# A Multi-Taxa Approach to Estuarine Biomonitoring: Assessing Vertebrate Biodiversity and Ecological Continuity using Environmental DNA Metabarcoding in the Rance River (Brittany, France)

**DOI:** 10.64898/2025.12.01.691502

**Authors:** Rachel Haderle, Alexandre Carpentier, Gael Kervarec, Anne Lize, Nils Teichert, Visotheary Ung, Jean-Luc Jung

**Affiliations:** ISYEB (MNHN) et station marine de Dinard; Universite de Rennes, BOREA (MNHN) et station marine de Dinard; Association Naturaliste d Ouessant; BOREA (MNHN), station marine de Dinard et School of Biosciences of Liverpool; BOREA (MNHN) et station marine de Dinard; ISYEB (MNHN)

**Keywords:** Environmental DNA (eDNA), estuarine biodiversity, metabarcoding: vertebrate diversity, ecological monitoring, Rance Estuary, tidal power plant, renewable marine energy

## Abstract

Estuaries are ecologically vital yet highly impacted ecosystems that serve as transitional zones between land and sea. Monitoring their biodiversity is essential but challenging due to their dynamic nature and the transient presence of many species. Traditionally actinopterygian monitoring in these systems still relies on conventional and intrusive methods such as gill nets and trawls. Environmental DNA (eDNA) metabarcoding offers a non-invasive, multi-taxa alternative that can complement these traditional approaches. Here, we applied an eDNA-based metabarcoding approach to characterize vertebrate diversity in the Rance Estuary, located in the Brittany Region of France. Water samples were collected from five stations spanning marine to freshwater environments. Special attention was given to two stations located upstream and downstream of the tidal power plant (TPP) dam to assess its potential impact on ecological continuity. We detected a total of 124 distinct vertebrate MOTUs—comprising actinopterygians, birds, mammals, and amphibians. Taxonomic composition followed the estuarine gradient, with Jaccard dissimilarity increasing with distance from the sea and largely driven by species turnover. While taxonomic and phylogenetic diversity remained relatively stable across the vertebrate community, functional diversity revealed an increasing terrestrial influence. For actinopterygians, taxonomic diversity decreased upstream, whereas phylogenetic and functional diversity indicated fine-scale structuring, even among nearby stations. This approach enabled the development of biodiversity metrics and facilitated comparisons with previous actinopterygian monitoring surveys in the same area based on conventional methods (scientific fishing using nets and dredges). Our results emphasise the potential of eDNA for holistic estuarine biomonitoring and establish a valuable baseline for future non-invasive assessments.

## Introduction

Estuaries are ecologically critical zones located at the interface between marine and freshwater environments, characterized by a diversity of physical, chemical, and topographic features (McLusky and Elliott, 2004). These areas typically harbor distinctive biodiversity, where spatial variations in salinity play a major role in species distribution (Gibson et al., 2023). Generally, species richness decreases from marine to brackish waters and then increases again in freshwater (Attrill, 2002). Estuaries typically host few resident species (Costanza et al., 1993) but play vital roles as habitats, nesting sites, and migration routes for many species (McLusky and Elliott, 2004). However, these ecosystems are subject to increasing anthropogenic pressures, including pollution and habitat degradation (Kennish, 2002). To preserve these ecosystems, conservation efforts are deployed, guided in Europe both by local regulations and area protective measures and by European environmental legislation such as the European Water Framework Directive (WFD) and the “Habitats” Directive (2000/60/CE and 92/43/CEE). In this global frame, actinopterygian monitoring programs have been developed across Europe, using various sampling methods, more or less invasive, which are sometimes difficult to implement, such as scientific fishing using nets and dredges, electrofishing, hydroacoustic, and Underwater Visual Census (UVC) techniques. Environmental genomics-based approaches analyzing environmental DNA (eDNA) now offer new possibilities for complementing these tried-and-tested methods with a non-invasive approach.

Environmental DNA (eDNA) originally refers to DNA extracted from an environmental sample without prior isolation of organisms (Birk et al., 2012; Coates et al., 2007; Delpech et al., 2010; Harrison and Kelly, 2013; Pérez-Domínguez et al., 2012; Taberlet et al., 2018). Biodiversity inventories can be captured from eDNA samples using a metabarcoding procedure, with the aim of assigning each eDNA molecule in the extract to its taxon of origin (Haderlé et al., 2024b; Jung, 2024; Valentini et al., 2009). eDNA metabarcoding is thus a non-invasive and powerful approach to study ecosystems that are difficult to sample and to detect rare or cryptic taxa (Bohmann et al., 2014; Günther et al., 2022; Haderlé et al., 2024a; Madon et al., In Press; Ruppert et al., 2019). Traditional biodiversity surveys and eDNA metabarcoding generally provide complementary insights (e.g. Rey et al. 2023).

Although most published eDNA studies have been conducted in freshwater environments (e.g. Bylemans et al., 2019; Deiner et al., 2016a; Jackman et al., 2021), and to a lesser extent in marine habitats (e.g. Albouy et al., 2015; Haderlé et al., 2024a; Rey et al., 2023), an increasing number of studies are now exploring the potential of eDNA metabarcoding to assess biodiversity in estuarine environments (Ahn et al., 2020; Cole et al., 2022; Gibson et al., 2023; Stoeckle et al., 2017). Most studies on vertebrates in estuaries have mainly concentrated on actinopterygians (Ahn et al., 2020; Zainal Abidin et al., 2022), very few implemented a broader and integrative approach that includes birds and mammals (Chiquillo et al., 2024; Ip et al., 2024; Saenz-Agudelo et al., 2022). Despite the recent development of eDNA integrative biomonitoring, this approach has proven effective. For example, in a subtropical estuary, eDNA metabarcoding outperformed trawling by detecting a greater diversity of pelagic and demersal actinopterygians, as well as elasmobranchs (Ip et al., 2024). In addition, eDNA has also proven useful for investigating habitat connectivity of fishes between marine and freshwater systems, as demonstrated in a Japanese river system by Yamanaka and Minamoto (2016).

Beyond generating biodiversity inventories, eDNA metabarcoding enables the exploration of multiple ecological dimensions—including taxonomic, phylogenetic, and functional diversity—thereby supporting more integrative ecosystem assessments. Although biodiversity inventory analysis—typically based on species richness and taxonomic identification—is essential, it is insufficient on its own to fully understand or mitigate biodiversity decline (Faith and Baker, 2006). A comprehensive approach must also incorporate components related to phylogenetic and functional diversity (e.g. Albouy et al., 2015; Safi et al., 2011). To our knowledge, while eDNA metabarcoding has been used to study vertebrate communities in estuaries and assess different aspects of biodiversity (Polanco et al., 2021), no one has simultaneously explored taxonomic, phylogenetic, and functional diversity to provide a comprehensive view of alpha and beta diversity patterns.

The Rance estuary in Brittany (France) is framed by two obstacles, an upstream dam with a lock limiting exchanges with the river section (“Le Châtelier” lock) and, downstream, a major dam formed by the tidal power plant (TPP) between Dinard and Saint-Malo, which influences connectivity between the river and the sea (Le Mao, 1986; Trancart et al., 2022). About 20 years after the TPP dam construction, Le Mao (1986) highlighted that the Rance estuary was an important spawning and nursery area for many species of actinopterygians, particularly flatfish, sole and plaice. Between 2012 and 2014, actinopterygian communities in the Rance estuary were surveyed using beam trawls as part of the Water Framework Directive (WFD). The ELFI (Estuarine and Lagoon Fish Index) indicated a poor ecological status, mainly due to low actinopterygian abundances compared to other French estuaries. In 2021, the AnaCoNoR project (Rault et al., 2023) conducted additional sampling using various techniques (plankton nets, beam trawls, fyke nets) to assess different actinopterygian life stage proportions. Results showed low ichthyoplankton densities and imbalances among ecological guilds (Rault et al., 2023). In a recent study, acoustic telemetry found that only one-third of tagged silver eel individuals reached the sea, suggesting the TPP dam may significantly hinder their migration (Trancart et al., 2022). In addition to ichthyofauna, the Rance estuary also serves as a critical habitat for various avian, mammalian, and amphibian. Every year, the estuary receives an influx of several thousands of wintering birds, highlighting its importance as a stopover and feeding site for migratory species (Retiere, 1994). Regarding mammals, the estuary supports both marine and terrestrial species. Notably, a humpback whale (*Megaptera novaeangliae*) was observed upstream of the TPP dam in February 2023, suggesting that under certain conditions – such as high tides, open sluice gates, or lock operations – large marine mammals can access inner parts of the estuary, despite the generally limited permeability of the structure. Semi-aquatic terrestrial species are also present in the estuary, taking advantage of its wetland environments.

This study applied vertebrate eDNA metabarcoding for a holistic, cross-sectional and multi-taxon approach to monitor biodiversity along an estuarine gradient. Its goal was to generate species lists from several stations positioned along the estuary, including two particular ones: one located outside the TPP dam on the marine side and the other one on the opposite side of the “Le Châtelier” lock (i.e. in freshwater). Based on these species’ lists, the alpha diversity was investigated at the taxonomic, phylogenetic, and functional levels across the major taxonomic groups detected. Beta diversities were also computed to analyse differences in taxonomic composition of different taxonomic groups (actinopterygians, mammals, birds, amphibians) between stations, with the primary objectives of assessing the effect that the TPP dam could have on the ecological continuity and determining whether the estuarine gradient is reflected in the detected taxonomic composition. In addition, this study offers the opportunity to compare eDNA metabarcoding results with those obtained from conventional monitoring techniques, such as the Water Framework Directive (WFD) assessments (Carpentier et al., 2014, 2013, 2012) and the AnaCoNoR survey (Rault et al., 2023). This comparative analysis provided valuable insights into the strengths and limitations of different methods for assessing vertebrate biodiversity in transitional aquatic ecosystems.

## Material and method

### 1. Sample collection and laboratory processing

The Rance ria is a relatively deep and narrow drowned river valley, over 20 km long, formed by the lower reaches of the Rance coastal river and which flows into the English Channel at the Bay of Saint-Malo (northwestern France). The ria is framed by two obstacles, a first dam with a lock at “Le Châtelier”, located upstream of the ria and which limits exchanges with the river, and a major lower dam supporting a tidal power plant (TPP) at the mouth of the basin.

Five sequential sampling stations were defined along the estuary (**Fig. 1**). Station 1 was located downstream of the estuary outside the TPP dam. Stations 2 to 4 were positioned progressively further from the sea, following the estuarine gradient. Station 5 was located in freshwater, upstream after the “Le Châtelier” lock.

**Fig. 1.**
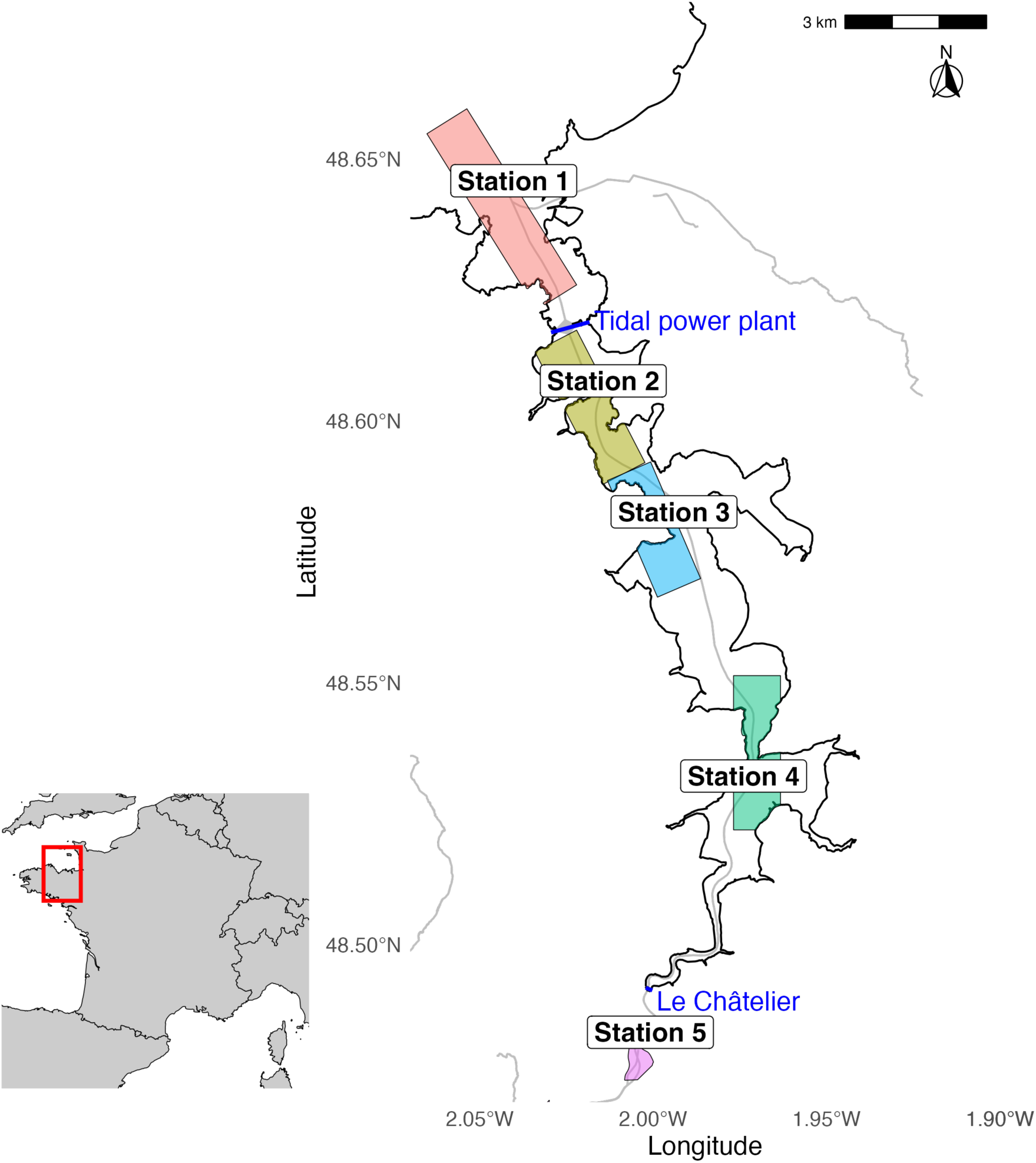
Map of the Rance estuary showing the five sampling transects (colored areas, from upstream to downstream). The upstream and downstream ends define each transect. The two dams (Rance tidal power plant and Le Châtelier) are indicated in blue.

In each station, water was continuously filtered, in duplicate on each side of a boat just below the surface, using an Athena peristaltic pump (Proactive, Hamilton, NJ, USA) through a VigiDNA 0.45 μm filtration capsule (SPYGEN, France) during 30 minutes at a ca. speed of 5 knots (Haderlé et al., 2024a). All samples were collected at high tide during two days in winter 2023 (the 2023-02-03 and the 2023-02-06) to focus on resident adult actinopterygian species, as the winter period does not align with the reproduction period of most species or the occurrence of larval and post-larval phases of actinopterygians in estuaries (Arevalo et al., 2023). Each capsule was then immediately filled with 80 mL of CL1 DNA preservation buffer (SPYGEN) and stored at room temperature until DNA extraction.

DNA extraction and amplification were performed by a dedicated DNA laboratory (SPYGEN, www.spygen.com). PCR amplification was performed using a universal vertebrate 12S mitochondrial rDNA primer pair Vert01 (5’ fwd-ACACCGCCCGTCACTCT, 5’ rev-CTTCCGGTACACTTACCATG; Taberlet et al. (2018)). The amplicons were sequenced using an Illumina MiSeq sequencer (Illumina, San Diego, CA, USA). The resulting sequence datasets (read sets) were analyzed using the OBITools package (Boyer et al., 2016).

Each MOTU, representing a grouping of sequences based on their molecular similarity, was associated with a number of reads per sample. The MOTUs were named according to the following nomenclature: Rance_n°MOTU; with Rance for the name of the estuary and a number corresponding to the order of appearance of the MOTU in the global list. To construct the biodiversity inventories, taxonomic assignments of each MOTU were meticulously checked by hand, considering molecular, taxonomic, and ecological criteria (Haderlé et al., 2024b). MOTUs served as the basis for deriving the list of detected taxa. In most cases, one MOTU corresponded to a single distinct taxon. However, two main exceptions were observed: (1) a single species could be identified from two or more MOTUs, and (2) several MOTUs could share the same list of potential species-level assignments, leading to more MOTUs than distinct taxa in those cases. In such situations, we inferred a single taxon from multiple MOTUs. For ecological information, local naturalist sources were consulted, including expert ornithologists, ichthyologists and the Atlas of Mammals of Brittany by the Groupe Mammalogique Breton.

Data are shared under a CC-BY 4.0 license on the GBIF: https://www.gbif.org/dataset/787386a7-aa69-4f99-ba95-9fd5a700705c. The publication of the data followed the recommendations outlined in the GBIF guide for DNA-derived occurrence data (Andersson et al., 2021), ensuring compliance with both the Darwin Core (DwC) framework (Wieczorek et al., 2012) and the MIMARKS standards (Yilmaz et al., 2011). Data standardization was performed following the metadata checklist and data formatting guidelines of Takahashi et al. (2025), using tools provided by the FAIRe suite, including FAIRe-ator, FAIRe-fier (Yong and Takahashi, 2025), and FAIRe2MDT for publication via the GBIF Metabarcoding Data Toolkit system (GBIF Secretariat, 2024).

All data processing and graphical representations were performed using R version 4.4.3 (R Core Team, 2025).

### 2. Comparisons of diversity values between sites along the estuarine gradient

The list of taxa identified at each station was first transformed into a binary presence/absence matrix (1: when the taxon was detected at the station, 0: when it was not). Analyses of phylogenetic and functional diversity were carried out for the 3 main taxonomic classes (actinopterygians, birds and mammals). To account for the bias in estimating diversity indices due to taxon detection linked to tributary inflows (DNA traces introduced into the estuary by freshwater inputs), strictly freshwater actinopterygian taxa detected in the estuary from Stations 1 to 4 were excluded from the calculations.

#### 2.1. Alpha Diversity

To compare the alpha diversity values specific to each site, taxonomic, functional, and phylogenetic diversity metrics were calculated.

##### 2.1.1. Taxonomic Alpha Diversity

The number of taxa from different taxonomic classes (species richness, SR) detected at each station were compared. To explore a potential correlation between the number of distinct taxa detected and the distance to the river mouth, normality was assessed using the Shapiro-Wilk test (with the shapiro.test function), and correlations were tested using the Pearson’s method (with the cor.test function).

Actinopterygians were classified into ecological guilds (Potter et al., 2015), widely employed in estuarine ecology (Elliott et al., 2007; Henriques et al., 2017). These ecological guilds distinguish estuary resident species (ER), freshwater species (FW), diadromous species (DIA) migrating through estuaries, juveniles of marine species (MJ) that use the estuary as a nursery, marine species (MA) sporadically present in estuaries, and marine straggler species (MS) that typically enter estuaries in low numbers and are most common in the lower reaches (Potter et al., 2015). Taxa that could not be assigned to a single category (approximately 25% of the actinopterygians) were excluded from the analysis.

##### 2.1.2. Phylogenetic Alpha Diversity

Phylogenetic diversity at each site was quantified using the PD index (Faith, 1992), which represents the sum of all branch lengths in the phylogenetic tree associated with the sampled taxonomic group. To compute this, we used the U.PhyloMaker package (Jin and Qian, 2023) to construct a comprehensive vertebrate megatree composed of a actinopterygian megatree (Rabosky et al., 2018), a bird megatree (Jetz et al., 2012), and a mammal megatree (Upham et al., 2019). All computations were performed using the picante package (Kembel et al., 2014). To include the influence of SR, PD values were standardized by computing Standardized Effect Size (SES) values (Gotelli and McCabe, 2002). SES values were calculated only when sufficient data was available. This was achieved by generating a null distribution of 1,000 randomized trees, in which species names were shuffled at the tips of the phylogenetic trees. Significant phylogenetic clustering or overdispersion was determined using the 95% percentile interval of a normalized Gaussian distribution.

##### 2.1.3. Functional Alpha Diversity

Functional alpha diversity was studied by computing the FRic index, corresponding to the volume of the convex hull polygon encompassing the species present in the synthetic functional space (Villéger et al., 2008), using the mFD package (Magneville et al., 2022).

Actinopterygian functional diversity was assessed using four categorical life-history traits that are widely employed to represent species’ habitat preferences and trophic roles in estuarine environments: body size, feeding habits, salinity tolerance, and vertical distribution within the water column. Trait data were sourced from Teichert et al. (2018b), who analyzed functional diversity patterns across 30 estuaries in France along the Northeast Atlantic coast. Body size was categorized into four ordered classes: ≤ 8 cm, 8 < and ≤ 15 cm, 15 < and ≤ 30 cm, and > 30 cm. Feeding habits were classified into six groups: piscivorous, omnivorous, planktivorous, herbivorous, benthic invertebrate feeders, and supra-benthic invertebrate feeders. Salinity tolerance was divided into four categories: marine, brackish, freshwater, and diadromous species. Lastly, vertical distribution in the water column was represented by three categories: pelagic, demersal, and benthic.

Similarly, functional traits were compiled for birds and mammals using the traitdata package and data from Wilman et al. (2014). Traits were selected to comply with those used for actinopterygian, ensuring comparability across taxonomic groups. For birds, these included body mass, dietary composition, aquatic habitat stratification, and vertical distribution, while for mammals, they encompassed body mass, dietary categories, and nocturnal activity patterns.

The FRic values were decorrelated from SR by computing Standardized Effect Size (SES; Gotelli and McCabe, 2002) values. SES values were obtained when sufficient data were available by subtracting the mean metric value across 1,000 random functional associations between species and their traits (null model) and dividing by the standard deviation of these null model metric values. SES values allowed the identification of sampling stations with functionally clustered or overdispersed taxa, regardless of SR. Assuming normality, SES values greater than 1.96 indicate significant functional overdispersion at a 5% significance level, while SES values below −1.96 indicate significant spatial clustering of taxa with specific traits (Leprieur et al., 2012).

#### 2.2. Beta-diversity

To identify the degree of uniqueness of each station in terms of taxonomic, phylogenetic, or functional diversity, we used a β-diversity approach. We considered the average values of taxonomic, phylogenetic, and functional β-diversity (and its two components: turnover and nestedness) for each pair of sampling stations. A principal component analysis (PCA) was conducted using the two components of each β-diversity index, employing the factoextra package (Kassambara and Mundt, 2017).

##### 2.2.1. Taxonomic Beta Diversity

Taxonomic β-diversity indices were computed using the beta.div.comp function from adespatial package (Dray et al., 2018) between the 17 stations using presence–absence Jaccard’s dissimilarity index and its two additive components: taxa turnover and taxa nestedness (Baselga, 2012). The first phenomenon, species replacement (or taxonomic turnover), occurs when a taxon present in sample A is absent from sample B, having been replaced by another species absent in sample A. The second phenomenon, nestedness, reflects differences in taxonomic richness between samples. In this case, sample B is a strict subset of sample A, containing only some of the taxa observed in A (Baselga, 2012). These two indices were related to the total biodiversity of the sample, highlighting the relative contributions of these factors to dissimilarity between samples (Dray et al., 2012).

##### 2.2.2. Phylogenetic Beta Diversity

At the regional scale (β-diversity), the UniFrac dissimilarity index was calculated, incorporating phylogenetic information and conceptually analogous to the taxonomic Jaccard index (Leprieur et al., 2012). This index ranges from 0 (all species in the two communities share the same phylogenetic history) to 1 (no shared phylogenetic history between species in the two communities). The UniFrac index was computed using the unifrac function from picante package (Kembel et al., 2014) and decomposed into two additive components: the UniFrac Turnover, quantifying the relative proportion of unique phylogenetic lineages between communities that is due to the replacement of unique evolutionary lineages, and the UniFrac Phylogenetic Diversity, which measures the amount of phylogenetic differences between phylogenetically nested communities (i.e., communities sharing at least one branch within a rooted phylogeny; Leprieur et al., 2012).

##### 2.2.3. Functional Beta Diversity

Functional β-diversity indices were estimated using the taxonomic-equivalent Jaccard dissimilarity index (Villéger et al., 2013) and computed with the beta.fd.multidim function from mFD package (Magneville et al., 2022). The functional β-diversity was decomposed into two additive components: the functional turnover, which refers to the replacement of functional strategies between communities (stations) within the multidimensional functional space, and the functional nestedness, defined as one community hosting a subset of the functional strategies present in the other (Villéger et al., 2013).

### 3. Comparisons of diversity metrics assessed on the Rance estuary between eDNA metabarcoding, the AnaCoNoR survey and the 2012-2014 WFD surveys

The aim of this analysis was to assess whether biodiversity metrics based on species richness, phylogenetic α-diversity, and functional diversity are comparable across surveys. These surveys differed both in sampling methods and spatial coverage within the estuary. To enable comparison, taxon lists from the three datasets were merged. The analysis specifically test whether these varying approaches yielded consistent diversity metrics. It is important to note that eDNA detections from station 5, located in a freshwater environment, were excluded, as were detections of several strictly freshwater species, which were assumed to result from river inflows. These exclusions were necessary because the two other surveys involved sampling exclusively within the estuary, without freshwater influence.

#### 3.1. WFD and AnaCoNoR surveys

The WFD surveys were conducted between 2012 and 2014 in the Rance Estuary. Sampling occurred annually during two seasons, spring and autumn, following a standardized protocol (Carpentier et al., 2014, 2013, 2012). The sampling method used was beam trawling. The corresponding dataset is publicly available at: https://surval.ifremer.fr/Donnees/Graphes-30140#/sensor/60005875. The primary objective was to provide an overall assessment of the water body’s status. Ecological condition was evaluated using a descriptive index of actinopterygian communities (ELFI). The results indicated that the ecological status of the Rance transitional water body was poor, with low actinopterygian abundances. This was particularly evident for diadromous species and juvenile marine actinopterygians, though estuarine resident species were less affected (Carpentier et al., 2014, 2013, 2012).

The objective of the AnaCoNoR survey was to characterize the functioning of the Rance catchment as a nursery area for juvenile actinopterygians, in relation to changes in hydro-sedimentary processes and the degree of connectivity with the marine environment. Scientific fishing campaigns were carried out in spring, early summer, and early autumn of 2021, using various sampling techniques, including plankton nets, beam trawls, and fyke nets (Rault et al., 2023). The ecological connectivity across the TPP dam was confirmed for the ichthyoplankton assemblage. Although the larval recruitment in the Rance estuary was relatively diversified, the ichthyoplankton density in the Rance estuary was much lower compared to the Bay of Saint-Malo, and imbalances were observed across ecological guilds (Rault et al., 2023).

#### 3.2. Diversity metrics

The species richness (SR), the phylogenetic diversity (PD), and their associated standardized effect size (SES) values were calculated, along with the functional richness (FRic) and its associated SES (see sections 2.1.2 and 2.1.3 for methods) for each dataset. To assess the significance of differences observed between the WFD, the AnaCoNoR, and the eDNA metabarcoding surveys, permutation tests were conducted for each biodiversity index. These tests involved 1,000 permutations, and the results were evaluated at a significant threshold of 0.05.

## Results

### 1. Alpha Diversity indices

A total of 124 distinct MOTUs was identified across all samples, corresponding to 118 detected taxa, including 66 actinopterygian MOTUs, 3 amphibian MOTUs, 36 bird MOTUs, and 19 mammalian MOTUs (**Table 1**). Among these, 74 were identified at the species level (**Table 1**). Actinopterygians were the main taxonomic class detected across the samples, from 43 taxa detected in station 1 to 17 in station 5. The diversity of actinopterygian taxa identified at each station decreased significantly with increasing distance from the river mouth (Pearson correlation test, cor = -0.97, p-value = 0.005) (**Fig. 2** and **Fig. 3**).

**Fig. 2.**
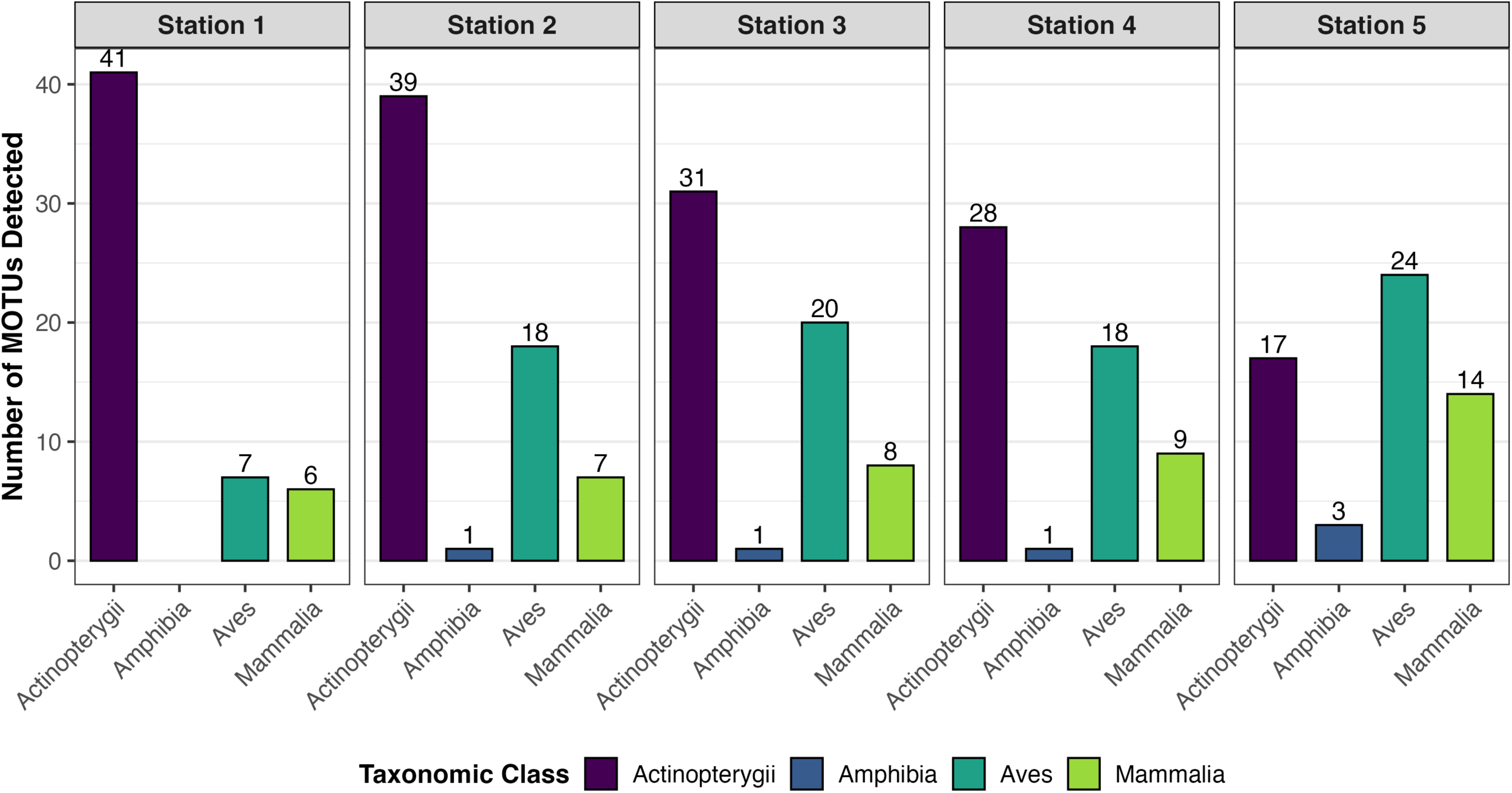
Number of MOTUs in each class detected for each station.

**Fig. 3.**
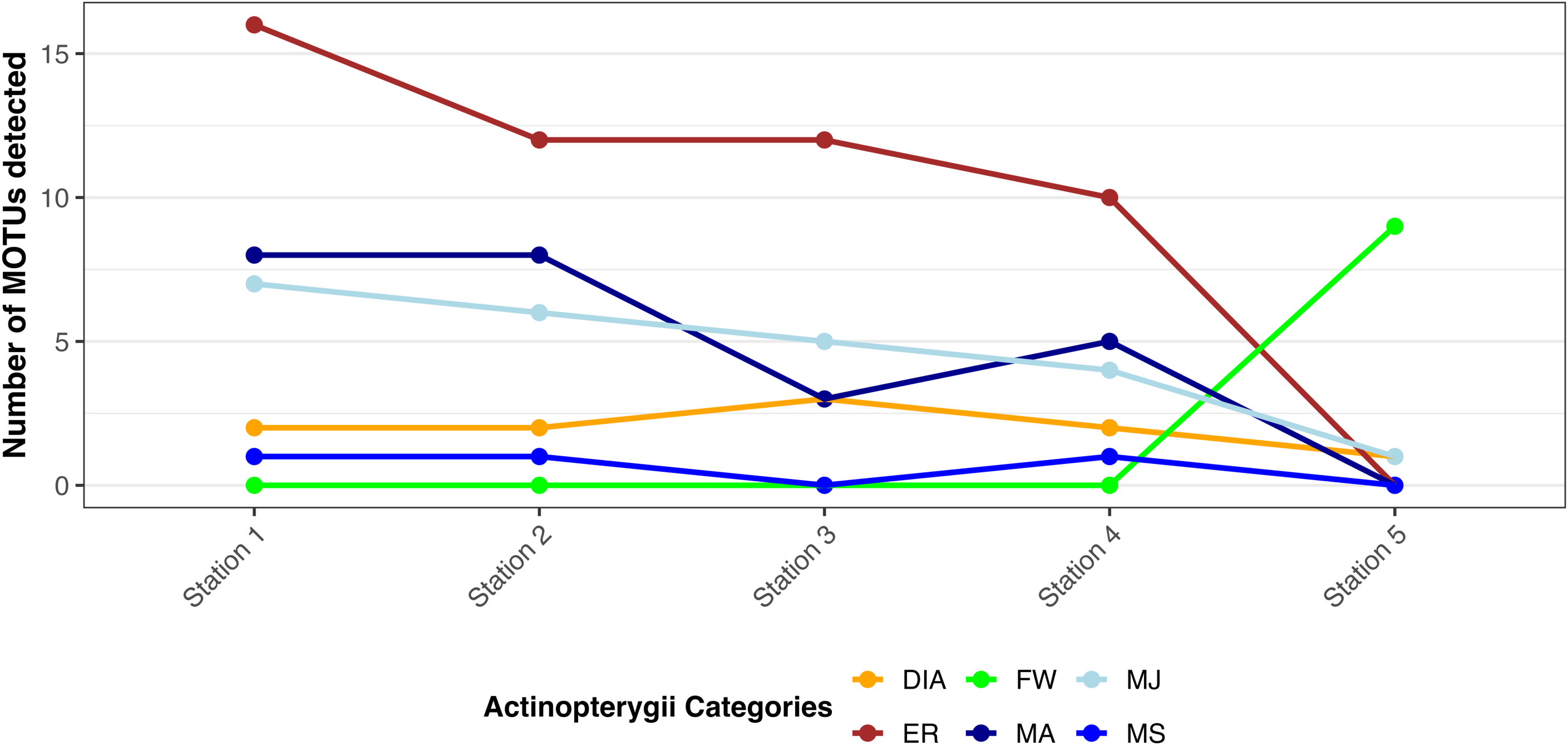
Number of actinopterygian MOTUs in each guild detected for each station.

**Table 1.**
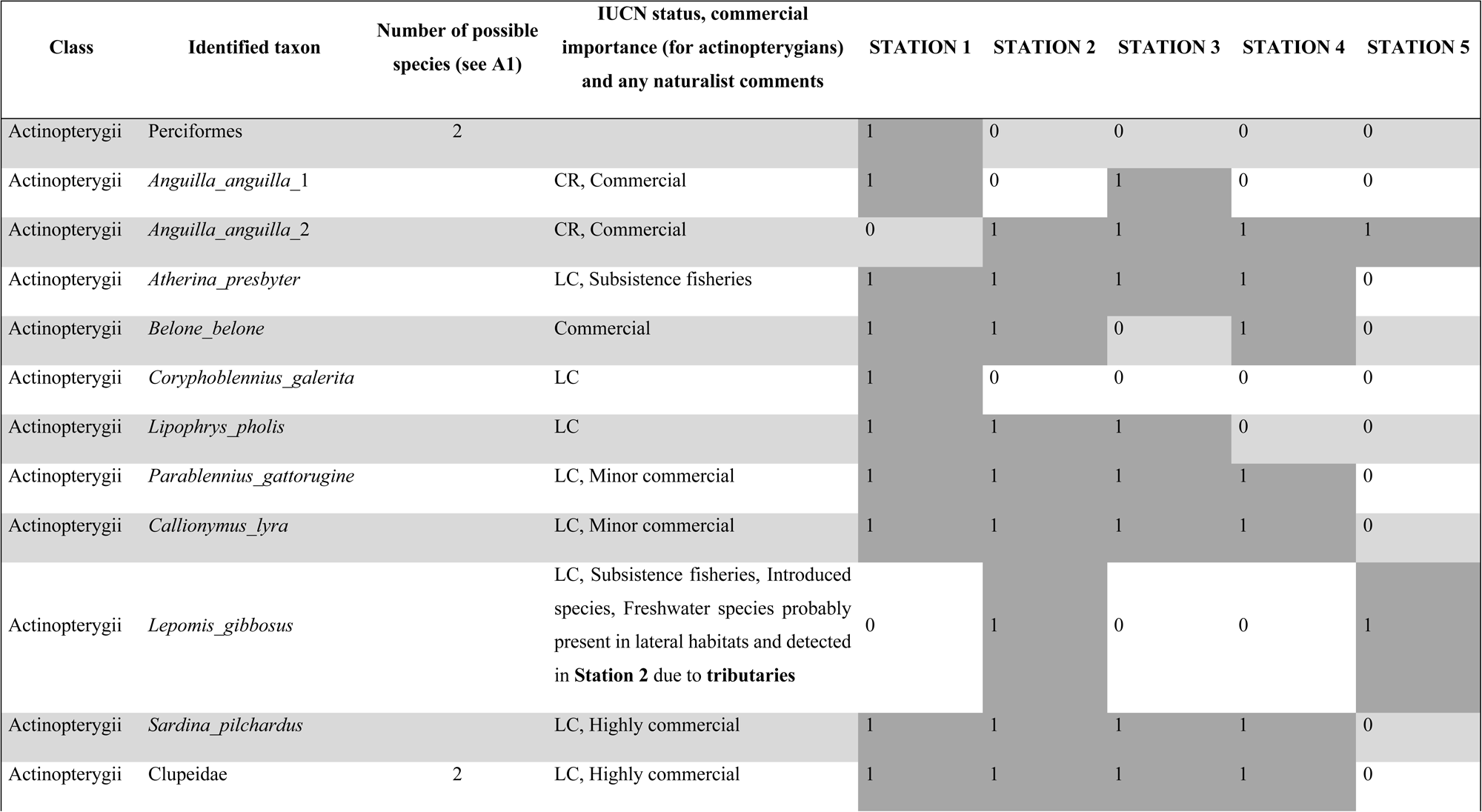
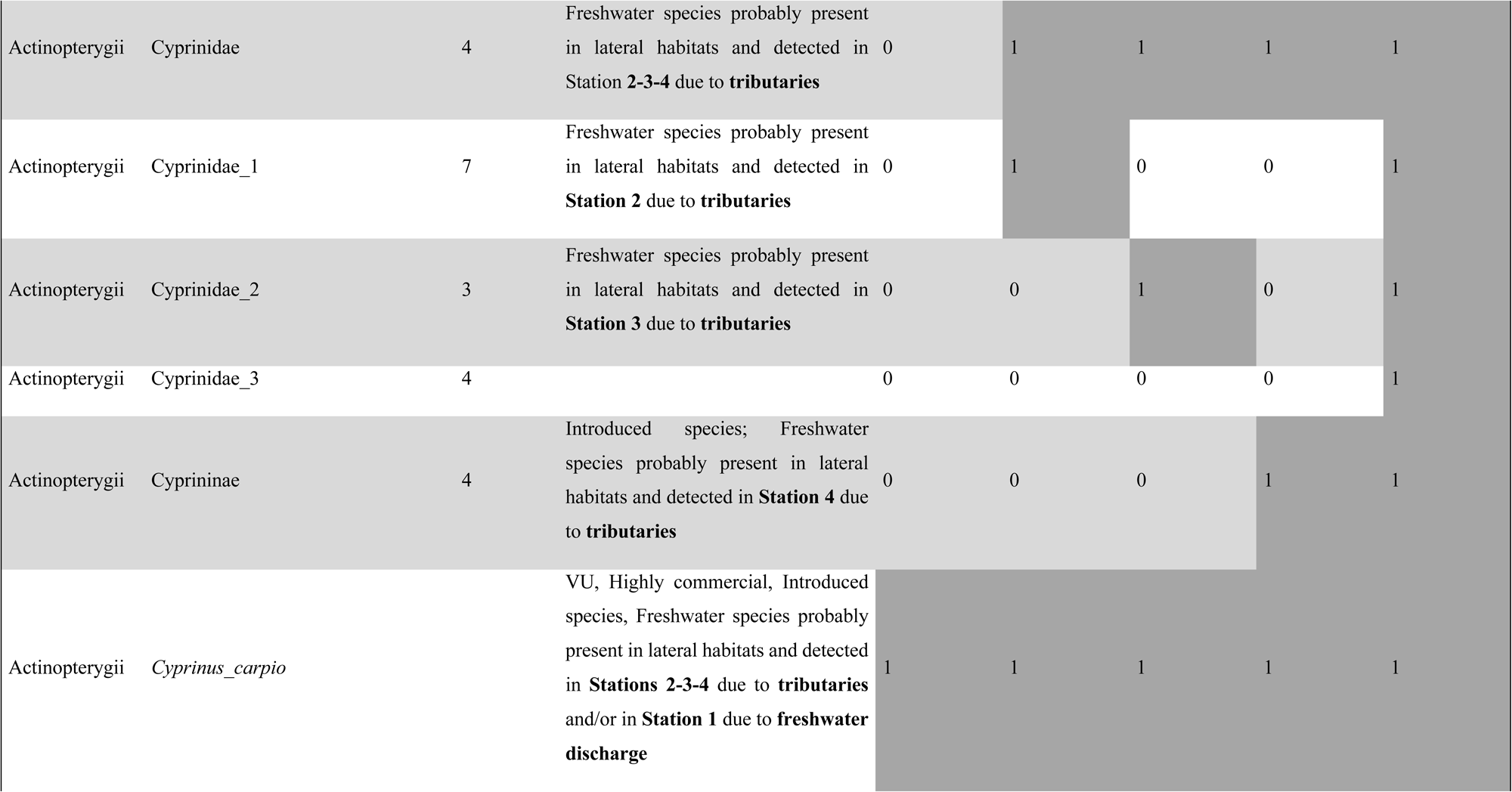
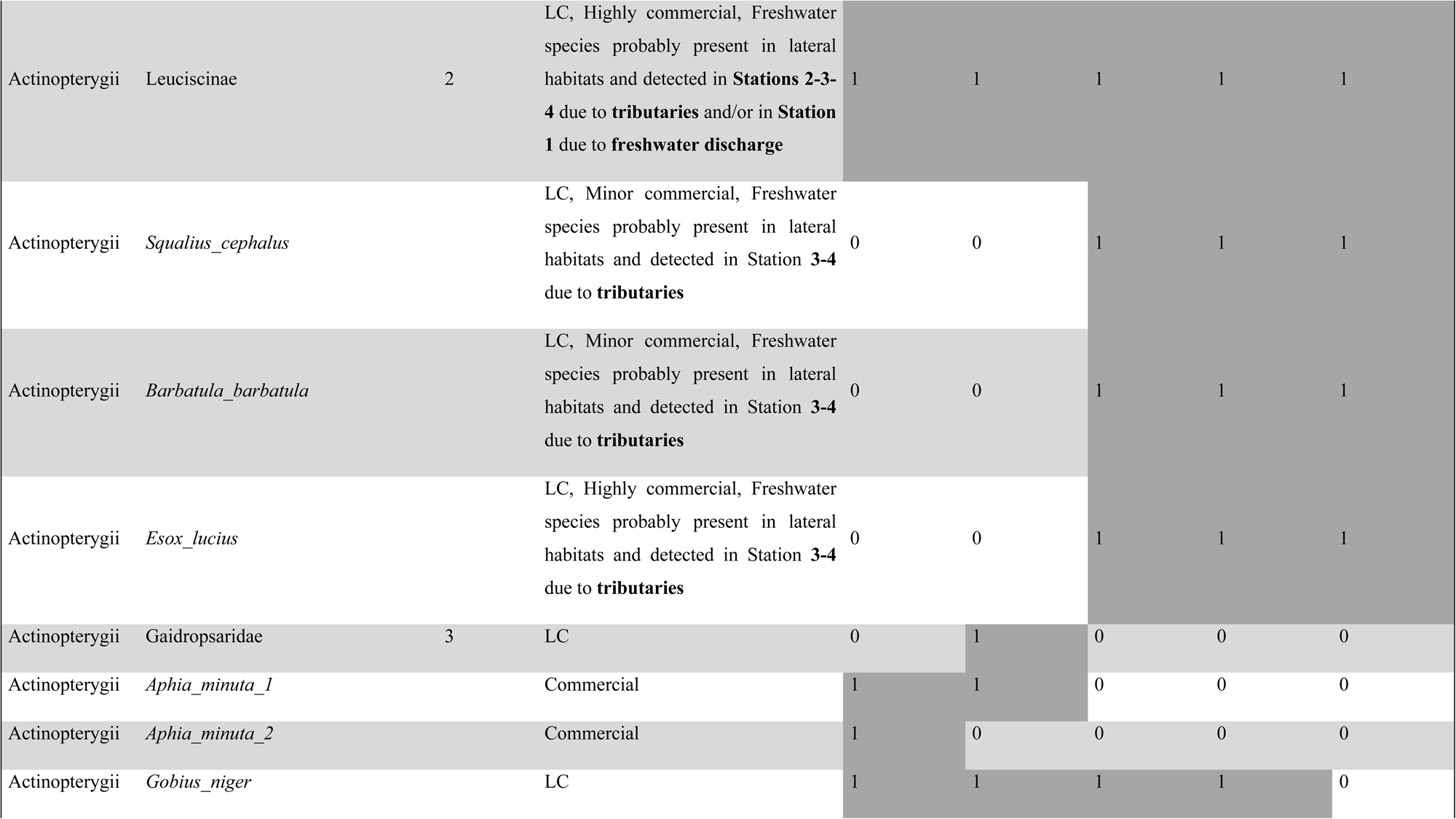
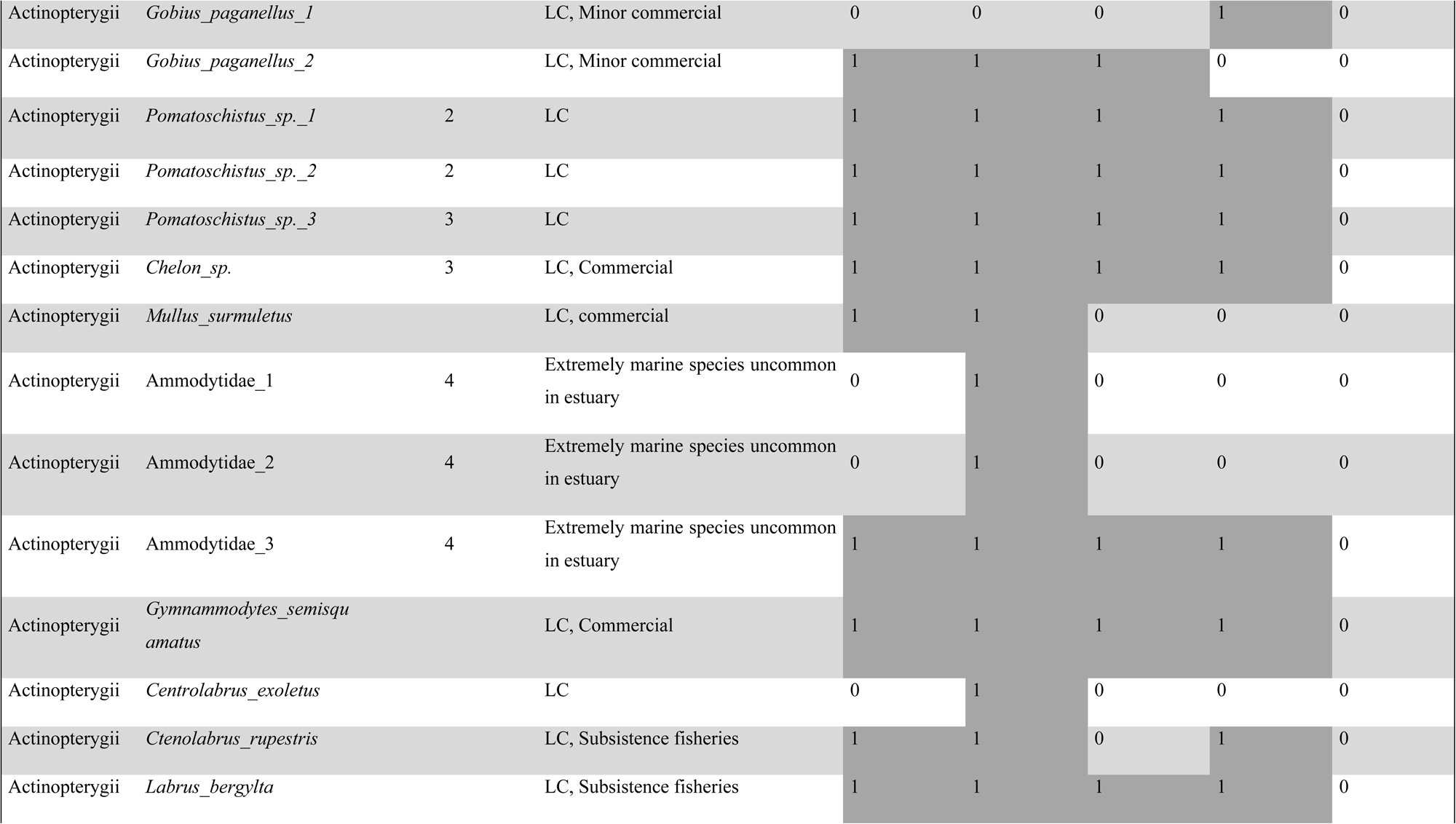
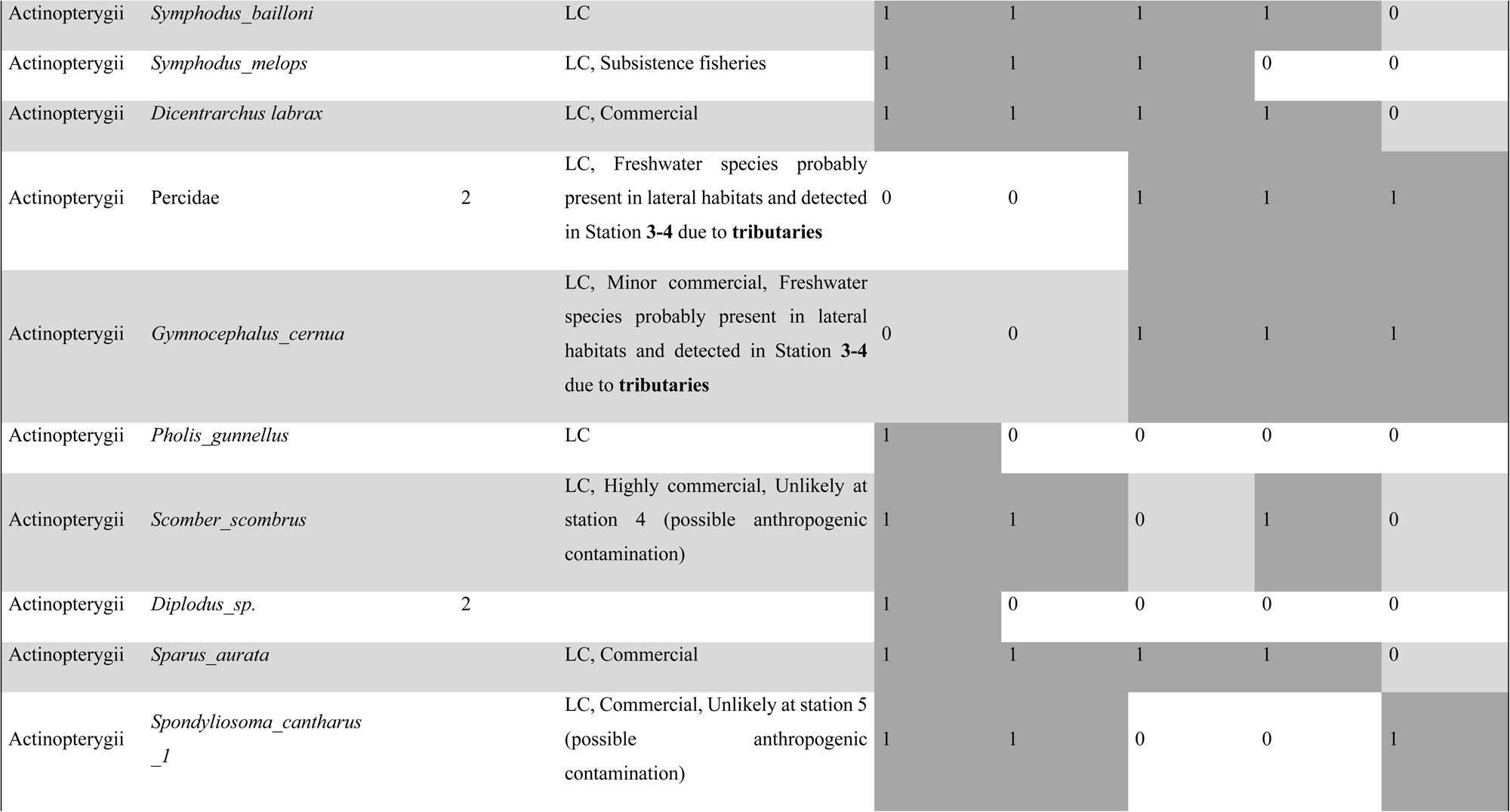
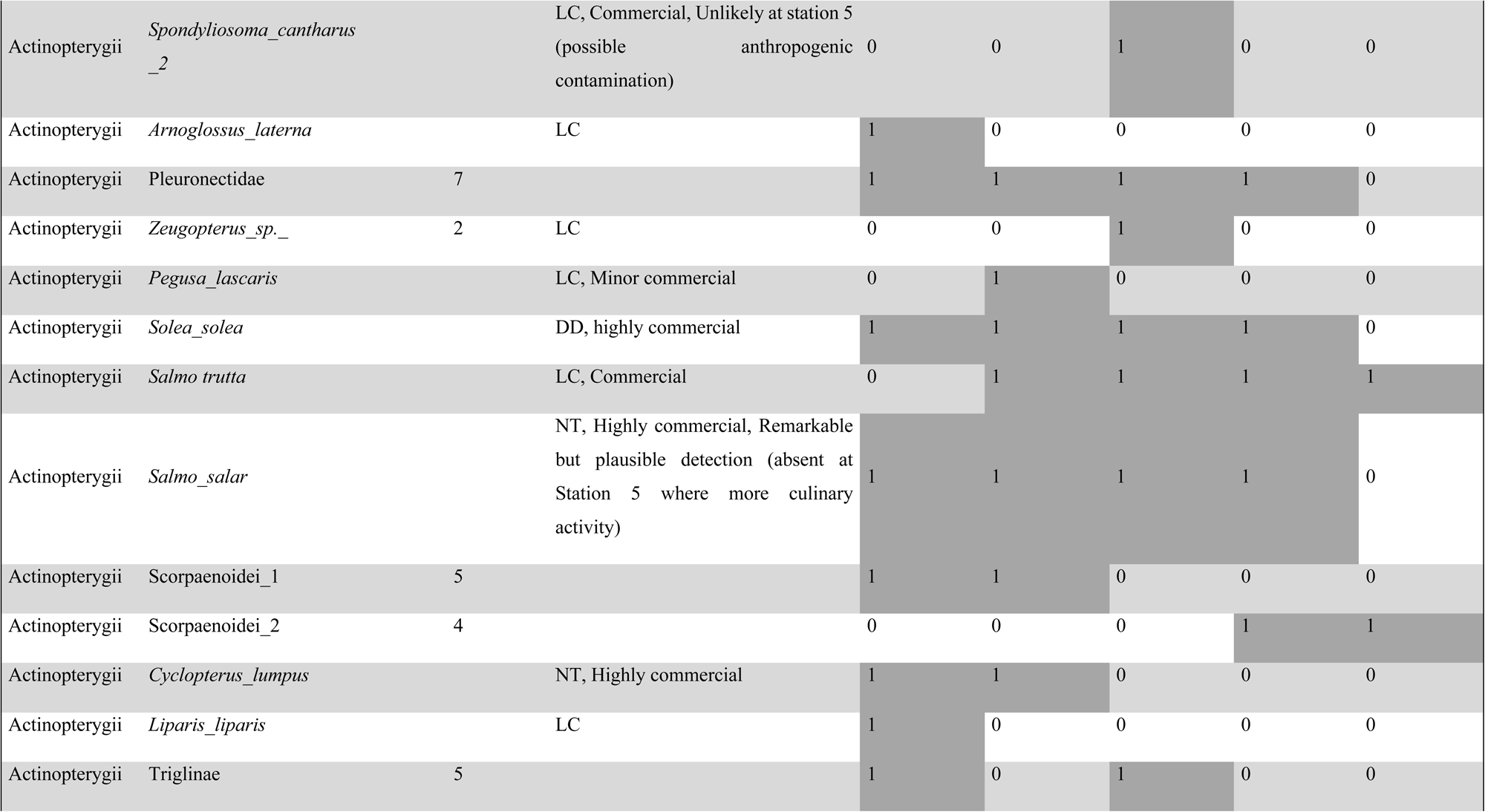
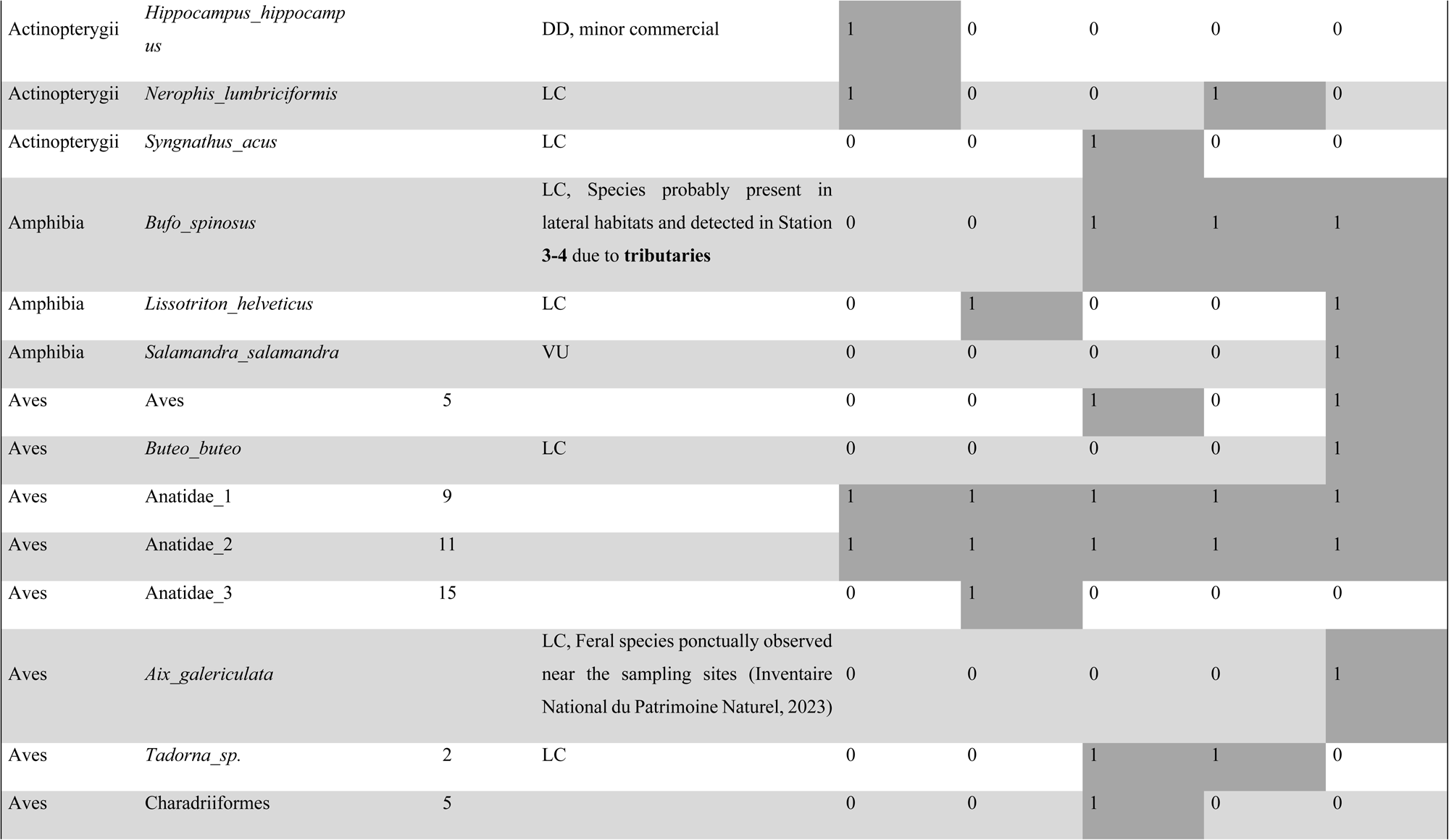
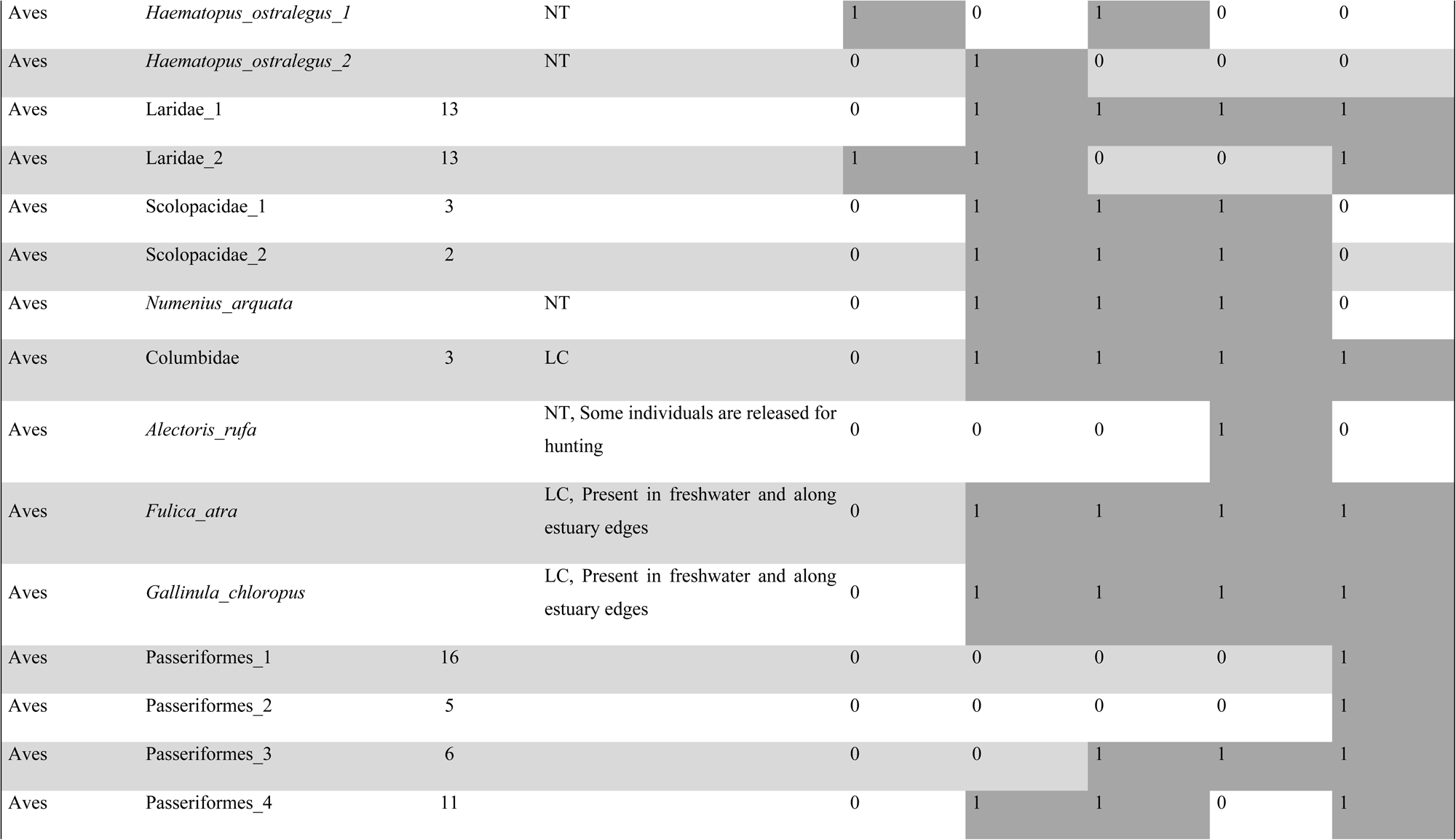
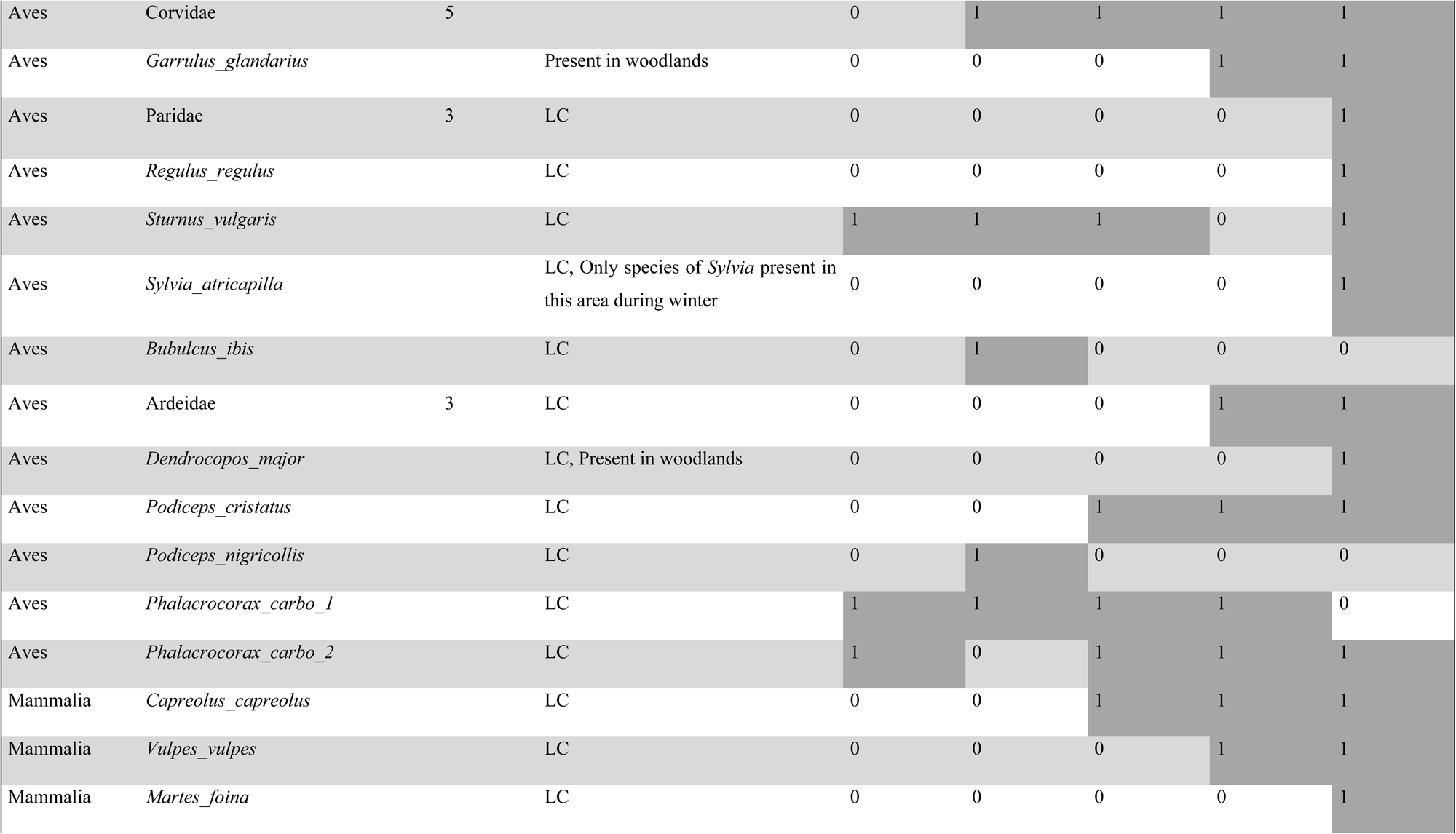
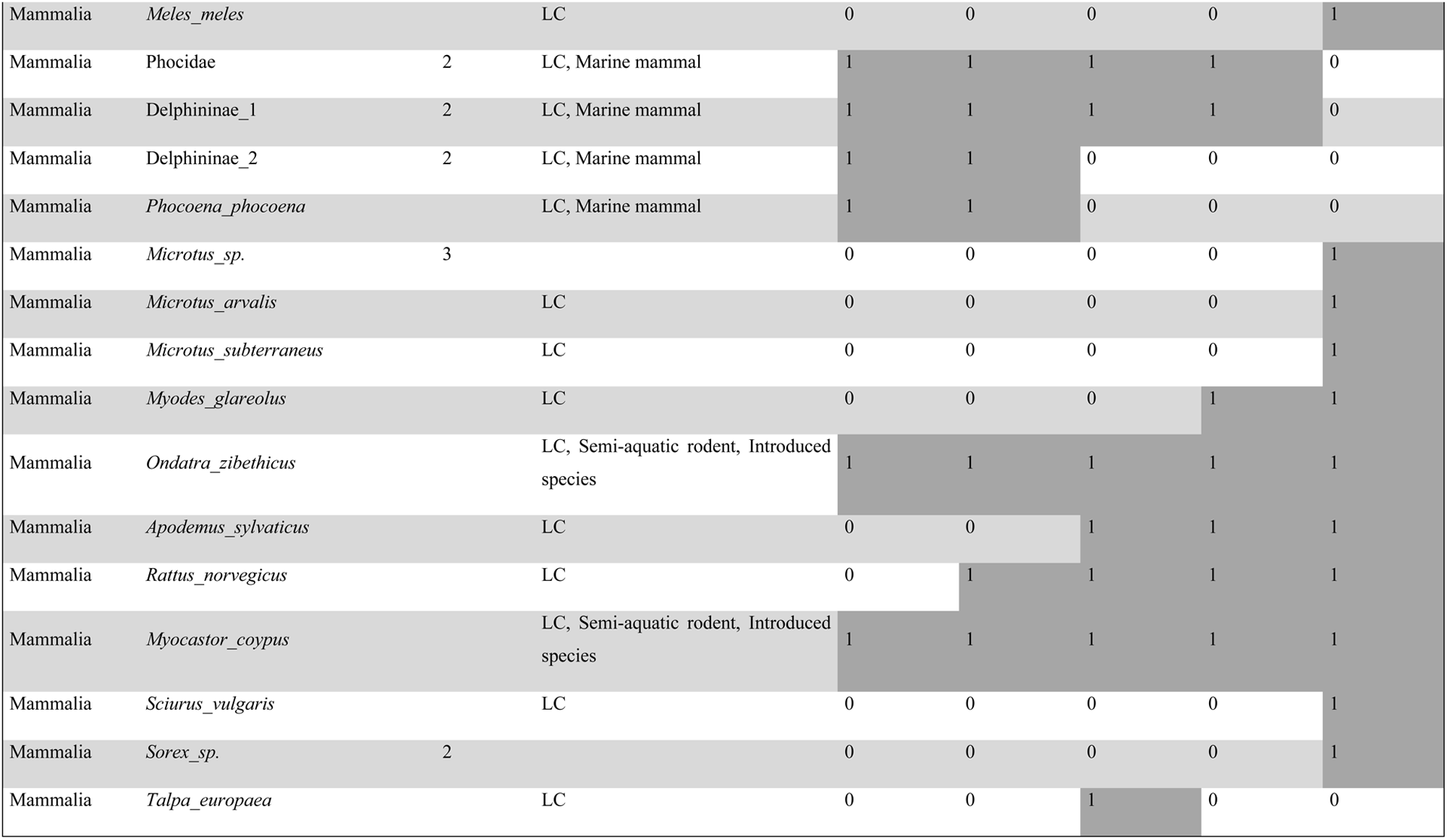
List of MOTUs and corresponding taxa detected at each station.

Several noteworthy species were detected (see **Table 1** and **Appendix A**), including introduced species and taxa listed on the IUCN Red List. Notably, the European eel (*Anguilla anguilla*), classified as critically endangered by the IUCN, was identified. Among the actinopterygians, several commercially important taxa were also detected. For birds and mammals, the detected species included both taxa closely associated with aquatic environments and others less specialized in such habitats.

Amphibians were only detected from station 2 onward (**Fig. 2**). Birds were detected at all stations, with a maximum of 24 detections at station 5 (**Fig. 2**). Mammals showed an increasing trend from stations 1 to 5 (**Fig. 2**), with a strong difference observed between the first four stations, where only marine mammals were detected (Phocidae, Delphininae, and *Phocoena phocoena*), and station 5, where no marine mammals were detected. Instead, station 5 revealed the presence of terrestrial mammals.

When separated by ecological guilds, estuary resident (ER) was the mostly represented category of actinopterygian, with between 16 (station 1) to 10 (station 4) ER taxa detected per station, albeit at station 5, where their numbers dropped to 0 (**Fig. 3**). Taxa associated with marine environments, such as juvenile marine species (MJ) and marine species (MA), exhibited similar trends to ER. Freshwater species (FW) were detected at all stations, with their numbers increasing as the stations were located upstream in the estuary. They accounted for most detections at station 5 (with 8 species out of the 10 detected).

#### 1.1. Alpha phylogenetic diversity (PD) values

The PD values for vertebrates (PD_vert) were relatively similar across stations, ranging from 4764 at station 1 to 6164 at station 4 (**Table 2**). For SES.PD_vert values, the lowest value was observed at station 1 (-0.97), followed by an increasing trend from stations 2 to 4, with a maximum at station 4 (1.70), significantly higher than expected (p-value = 0.952) given the taxonomic richness. Subsequently, a decrease was noted at station 5, reaching a value above station 1 (0.16).

**Table 2.** Phylogenetic diversity values calculated for all vertebrates (PD_vert) and for actinopterygians only (PD_actino), along with their associated standardized effect sizes (SES.PD_vert and SES.PD_actino); asterisks next to SES values indicate statistical significance (p > 0.95).

|  | STATION 1 | STATION 2 | STATION 3 | STATION 4 | STATION 5 |
| --- | --- | --- | --- | --- | --- |
| PD_vert | 4764 | 6144 | 5871 | 6164 | 5483 |
| SES.PD_vert | -0.97 | 0.45 | 1.10 | 1.70* | 0.16 |
| PD_actino | 2830 | 2566 | 2696 | 2644 | 1268 |
| SES.PD_actino | 0.82 | -0.59 | 1.16 | 2.21* | -1.45 |

The PD values for actinopterygians only (PD_actino) ranged from 2830 at station 1 to 1268 at station 5. Station 1 exhibited the highest PD_actino value, while stations 2 through 4 displayed relatively similar values. The lowest PD_actino value was observed at the freshwater station 5. The lowest SES.PD_actinopterygian value was also recorded at station 5 (-1.45), while the highest was observed at station 4 (2.21), which was significantly higher than expected (p-value = 0.995) given the taxonomic richness.

#### 1.2. Alpha functional diversity values

Functional richness values (FRic) were calculated for each major taxonomic group detected (actinopterygians, birds, and mammals). The missing values (NAs) correspond to stations where insufficient data were available. SES.FRic values were calculated only for actinopterygians due to the lack of sufficient data for the other groups.

The FRic_actino values were relatively consistent across the first four stations (**Table 3**), with station 4 exhibiting the highest value (22e-04) and station 3 the lowest (6.1e-04). Freshwater station 5 displayed the lowest FRic_actino value overall (1.3e-04), more than 15 times lower than the station 4. The SES.FRic_actino values showed significant spatial clustering of species sharing similar traits at each station. Like the FRic_actino values, SES.FRic_actino values were highest at station 4 (-2.45), relatively consistent across stations 1 to 3 (average around -3.5), and lowest at station 5 (-4.24).

**Table 3.** Functional diversity values calculated for actinopterygians (FRic_actino), birds (FRic_birds) and mammals (FRic_mamm), along with their associated standardized effect sizes when enough data; asterisks next to SES values indicate significant spatial clustering.

|  | STATION 1 | STATION 2 | STATION 3 | STATION 4 | STATION 5 |
| --- | --- | --- | --- | --- | --- |
| *FRic_actino* | 14<sup>e</sup>-04 | 10<sup>e</sup>-04 | 6.1<sup>e</sup>-04 | 22<sup>e</sup>-04 | 1.3<sup>e</sup>-04 |
| *SES.FRic_actino* | -3.17* | -3.65* | -3.72* | -2.45* | -4.24* |
| *FRic_birds* | NA | 2.5e-04 | 5.5e-04 | 2.7e-04 | 11e-04 |
| *FRic_mamm* | NA | NA | 1.3e-04 | 0.9e-04 | 6.2e-04 |

FRic_birds and FRic_mammals showed a similar pattern, with relatively low values observed at stations 2 to 4, and the highest values recorded at station 5 (insufficient data were available for station 1 for birds and mammals, and station 2 for mammals).

#### 1.3. Benchmarking of Taxonomic, Phylogenetic, and Functional Actinopterygian Alpha Diversities

The taxonomic richness, SES of phylogenetic diversity (PD), and SES of functional diversity (FRic) values for actinopterygian were normalized across all stations and represented on radar charts (**Fig. 4**). Station 1 exhibited the highest taxonomic richness, along with strong phylogenetic and functional diversity values. In contrast, station 5 showed low values for all three diversity indices. Stations 2 and 3 displayed intermediate values for the three metrics. Notably, station 4 demonstrated the highest phylogenetic and functional diversity values, despite having the second-lowest taxonomic richness.

**Fig. 4.**
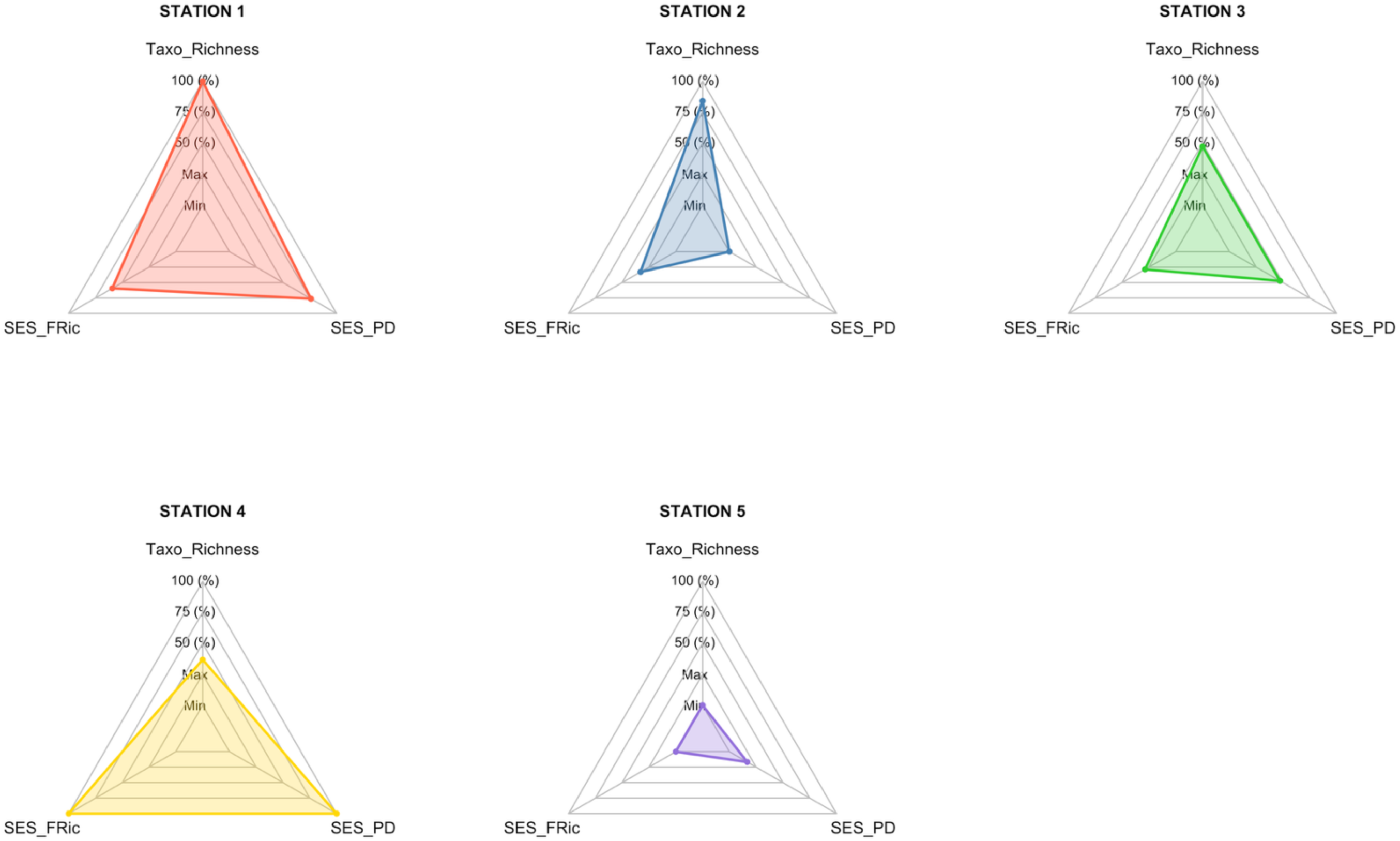
Radar charts showing, for each station, the standardized values (scaled between 0 and 1) of taxonomic richness (Taxo_Richness), the standardized effect size of functional richness (SES_FRic), and the standardized effect size of phylogenetic diversity (SES_PD) for actinopterygian taxa.

### 2. Results of beta Diversity indices

For actinopterygians, Jaccard dissimilarity values increased with distance from the river mouth, ranging from 0.40 between stations 1 and 2 to 0.98 between Stations 1 and 5. Overall, taxonomic turnover, accounting for 63% of the variation, was the primary driver of actinopterygian taxonomic dissimilarity, followed by nestedness, explaining 37% of the variation. Similar patterns were observed for actinopterygian functional beta diversity, with 71% of variation attributed to turnover and 29% to nestedness, and for phylogenetic beta diversity, with 76% for turnover and 24% for nestedness. These results emphasize the contribution of both components, with turnover being the predominant factor across all actinopterygian diversity facets.

A similar pattern was observed for mammals, with Jaccard dissimilarity values ranging from 0.14 between stations 1 and 2 to 0.89 between stations 1 and 5. Taxonomic dissimilarity was mainly driven by turnover (56%), with nestedness accounting for 44%. Variation of functional beta diversity was more strongly influenced by turnover (78%) than nestedness (21%), indicating that turnover played a more dominant role in shaping functional diversity differences. Phylogenetic beta diversity was more balanced, with 51% of its variation explained by turnover and 48% by nestedness.

For birds, the pattern was slightly different. Jaccard dissimilarity indices ranged from 0.81 between stations 1 and 4 and between stations 1 and 5, to 0.35 between stations 3 and 4. The taxonomic dissimilarity was nearly equally partitioned, with turnover explaining 52% and nestedness 48% of its variation. However, the variation of the functional beta diversity was primarily influenced by nestedness (83%) rather than turnover (17%), indicating that differences in functional diversity were largely driven by species loss rather than species replacement. In contrast, the variation of the phylogenetic beta diversity was more influenced by turnover (63%) than by nestedness (37%).

A Principal Component Analysis (PCA) was performed to visualize the spatial arrangement of stations based on their taxonomic, functional, and phylogenetic diversity components (**Fig. 5**) for actinopterygians, birds, and mammals. Actinopterygians and mammals followed a similar trend, with the horizontal axis being the most discriminant, primarily driven by taxonomic and phylogenetic turnover, which strongly distinguished station 5. Station 1 was mainly discriminated along the vertical axis, while stations 2, 3, and 4 appeared relatively close. In contrast, for birds, station 1 appeared strongly different than other stations, primarily due to the influence of phylogenetic and taxonomic nestedness.

**Fig. 5.**
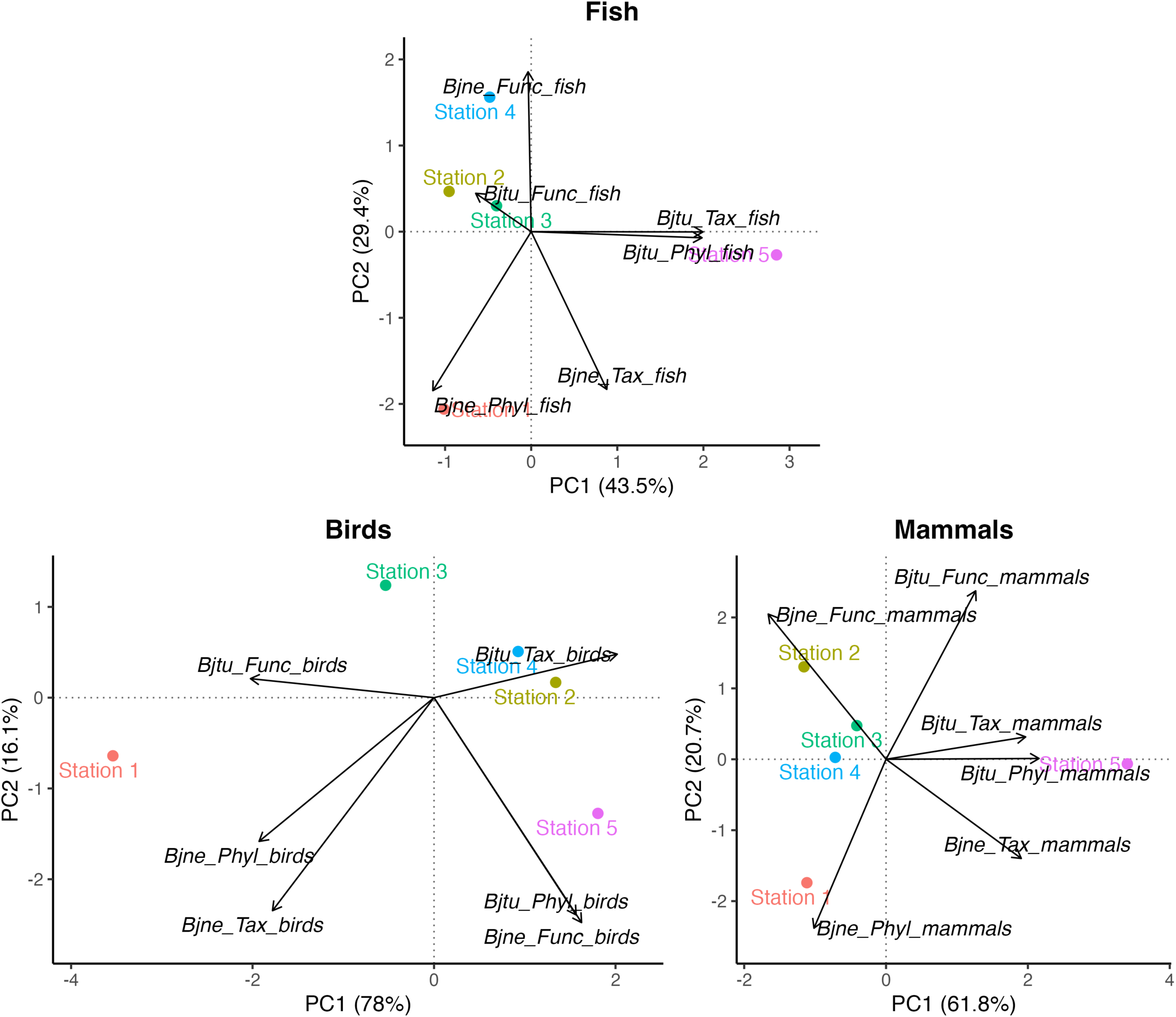
Principal Component Analyses (PCA) for actinopterygians (top), birds (bottom left), and mammals (bottom right), showing the spatial arrangement of stations based on calculated beta diversity values. Variables indicate the contribution of turnover (Bjtu) and nestedness (Bjne) components to taxonomic (Tax), phylogenetic (Phyl), and functional beta diversity (Func).

### 3. Inter-approach Comparison of Diversity Metrics (eDNA-, beam trawling- and ichtyoplancton-based analysis)

The diversity metrics calculated across the three surveys (WFD, based on beam trawling, Anaconor, based on ichtyoplancton identification, and this study based on eDNA metabarcoding) were generally similar (**Fig. 6**), although some subtle despite minor differences were observed. For species richness (SR), eDNA showed slightly higher values (53) compared to AnaCoNoR (48) and WFD (45), although all values were in the same range. In terms of functional diversity, eDNA values were intermediate between WFD (21e-04) and AnaCoNoR (41e-04), with SES.FRic values showing a similar trend.

**Fig. 6.**
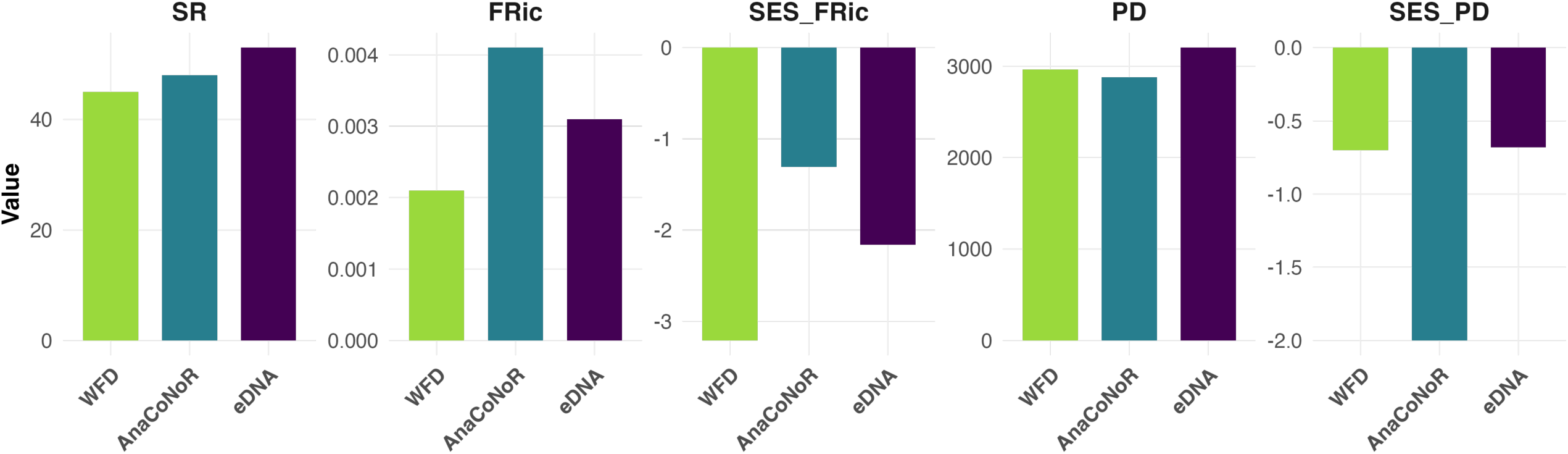
Graphs of values of species richness (SR), functional diversity (FRic) and phylogenetic diversity (PD) along with their associated SES values, calculated for the estuarine actinopterygian community along the Rance estuary using three methods: WFD surveys, AnaCoNoR, and eDNA (this study).

For phylogenetic diversity (PD), eDNA exhibited slightly higher values (3203) compared to AnaCoNoR (2879) and WFD (2964), which could likely be attributed to the higher SR observed with eDNA (**Fig. 6**). The SES.PD values showed that eDNA results were similar to those obtained from WFD, whereas AnaCoNoR exhibited a distinct pattern, with SES values significantly lower than expected based on the observed SR, indicating a divergence from the expected phylogenetic patterns.

A permutation test (n=1,000) revealed no significant differences in the calculated indices between the three datasets (**Appendix B**), further confirming the overall homogeneity of the diversity metrics across the methods.

## Discussion

The aim of this study was to assess how eDNA metabarcoding captures vertebrate species diversity in an estuarine system, in comparison with the known local diversity established through a range of traditional, and generally more invasive, survey techniques. Alpha and beta taxonomic, phylogenetic, and functional diversity metrics were calculated notably regarding the estuarine salinity gradient. Additionally, the ecological continuity of detected taxa on either side of the TPP dam was examined. The results were compared with actinopterygian species lists from previous studies using different methodologies (ichtyoplancton analysis and beam trawling sampling), to assess whether various diversity metrics reveal consistent patterns.

### 1. Naturalist Inventory and Taxa Detection

For actinopterygians, 11 species, excluding strictly freshwater taxa, were detected exclusively with eDNA metabarcoding, these species were undetected by classic actinopterygian surveys in the 2000s. Some of these species had already been identified in Le Mao’s (1986) study, such as *Cyclopterus lumpus*, *Pholis gunnellus*, *Sardina pilchardus*, *Scomber scombrus*, and *Sparus aurata*. Their detection confirms the capacity of eDNA metabarcoding to identify taxa that are less abundant or more difficult to sample using traditional survey methods (e.g. Bohmann et al., 2014; Valentini et al., 2016). This corroborates previous findings that eDNA is a powerful tool for investigating ecosystems that are challenging to sample and for detecting rare or cryptic taxa (Bohmann et al., 2014; Ruppert et al., 2019). For instance, *Phocoena phocoena* (the harbour porpoise) was detected at both station 1 and station 2, despite its cryptic behaviour (i.e. visual observations are difficult). eDNA allowed reliable detection, consistent with previous results in the Iroise Sea during summer 2019, when the species was repeatedly detected despite limited visual records (Jung and Gicquel, 2021). Interestingly, signals of *Salmo salar* were detected at stations 1 to 4, although this diadromous species is temporarily passing in estuaries to reproduce in freshwater (McCormick et al., 1998). However, because of its importance in human food these detections must be interpreted with caution, as they can result from anthropogenic sources rather than *in situ* presence.

Marine and freshwater species represent high proportions of assemblages in estuaries but are unlikely to co-occur in the same habitat within an estuary due to their divergent salinity tolerances (i.e., marine species typically occur downstream and freshwater species upstream; Whitfield et al., 2012). The detection of strictly freshwater species at stations 1-4 suggests the possible influence of freshwater tributaries, as these species cannot survive in estuarine conditions because of osmotic constraints (e.g. Whitfield, 2015). Such detections are possible in connected systems, where tributaries or lateral freshwater habitats may act as temporary refuges or sources of DNA, as shown in both Euclide et al. (2021) and Pavkovic et al. (2025).

This study showed a decrease in the number of actinopterygian taxa going upstream along the estuary. This result aligns with the conceptual model of estuarine biodiversity changes along the salinity continuum proposed by Whitfield et al. (2012). The low taxonomic richness observed at station 5 may be related to the fact that this site corresponds to the beginning of the freshwater zone, physically separated from marine waters by Le Châtelier lock. Station 5 also stands out as the most distinct in terms of taxonomic composition. It is likely that the number of freshwater actinopterygian taxa would have increased if additional upstream stations had been included in the sampling.

As shown by Polanco et al. (2021), river eDNA conveys both aquatic and terrestrial environmental signals, and thus serves as an integrator of biodiversity information. In addition to actinopterygians, eDNA metabarcoding can also detect amphibians, birds typical of these habitats, such as *Phalacrocorax carbo*, as well as species typically found in woodlands, whose DNA traces are found in the water (e.g. *Dendrocopos major* at station 5). Terrestrial vertebrates are well detected through metabarcoding of aquatic eDNA in systems close to terrestrial environments (e.g. Harper et al., 2019; Zhang et al., 2023). This is similar for mammals, and in addition to marine mammals known to cross the Rance TPP dam, like the common dolphin (*Delphinus delphis*) or the bottlenose dolphin (*Tursiops truncatus*), and to semi-aquatic mammals (*Myocastor coypus* and *Ondatra zibethicus),* species which are typically associated with terrestrial environments, like *Capreolus capreolus*, were also detected.

### 2. An Estuarine Gradient that does not appear to be impeded by the TPP Dam, as Evidenced by Taxonomic Composition and Diversity Values

In estuaries, species distribution is highly constrained and structured by fast-changing abiotic conditions and ecological interactions (Henriques et al., 2017). The list of vertebrates detected by eDNA revealed a gradual replacement of marine species by freshwater species, along with the presence of some terrestrial taxa such as birds and mammals. This pattern was particularly clear for actinopterygians, about 56% on average of the detected taxa. Their composition shifted progressively along the estuarine gradient, reflecting the transition from saline to freshwater conditions. Filtering for freshwater species, presumably detected due to the tributaries, allows us to focus on the estuarine horizontal gradient rather than the lateral one, which is also known to structure the taxonomic and functional assemblages of actinopterygians (Pavkovic et al., 2025).

The potential downstream homogenization effect due to eDNA transport from upstream is often considered a major challenge when using eDNA metabarcoding in lotic ecosystems (e.g. Deiner et al., 2016; Deiner and Altermatt, 2014; Jane et al., 2015; Laporte et al., 2021). In this study, the clear presence of an estuarine gradient in taxonomic composition suggested that eDNA variation in the Rance is horizontally structured and not strongly influenced by eDNA passive transport from the Rance river. Similar findings have been reported in other studies, such as in a river system in Switzerland (Mächler et al., 2019) and in the Gulf of St. Lawrence (García-Machado et al., 2022), where taxonomic turnover between sites was observed, further suggesting that eDNA transport is not sufficient to obscure the signatures of community changes.

Ongoing environmental changes in estuarine ecosystems place significant physiological stress on resident organisms, restricting the range of species that can successfully establish themselves (Elliott and Quintino, 2007). Focusing on actinopterygians, diversity values were lowest at the freshwater station, although functional and phylogenetic diversities were decorrelated from taxonomic richness. The marine station showed the highest values across all three metrics. A decline in diversity indices was observed between station 1 and stations 2 and 3. At station 4, we observed high functional and phylogenetic diversity values, as well as significant standardized effect sizes, despite relatively low taxonomic richness. This station is in a zone where salinity remains relatively stable (around 32.5 at high tide; Rault et al. 2023), indicating marine-like conditions. This pattern may reflect a paradox: although salinity shows little variation up to station 4, a significant shift in actinopterygian community structure was observed. Therefore, factors beyond immediate salinity differences—such as slight environmental gradients or upstream transitions (notably between stations 4 and 5)—may play a key role in shaping community assemblage in this part of the estuary. Overall, the balance between functional and phylogenetic diversity across stations may reflect niche conservatism—the tendency of closely related species to share ecological traits (Wiens et al., 2010; Wiens and Graham, 2005). However, because phylogenetic diversity does not always reflect functional differences (Mazel et al., 2018), it is important to examine both aspects independently to gain a clearer understanding of community structure and ecosystem function.

The phylogenetic diversity of all vertebrates’ values was very similar across stations, indicating relative stability, athough they were slightly lower at station 1. This can be explained by the predominance of actinopterygian species and the rare presence of birds and mammals, which are phylogenetically more distant than the detected actinopterygian species. The functional diversities of birds and mammals’ values were highest at station 5, likely due to an increase in terrestrial species detections as one moved further inland. However, this pattern could not be disentangled from species richness because of the limited number of detections.

Although the number of taxa detected was similar between stations, strong dissimilarities were observed between pairs of stations. Beta diversity calculations revealed that taxonomic dissimilarities of actinopterygians and mammals increased with the distance between stations, with the greatest difference between stations 1 and 5. For all three components of beta diversity of actinopterygian and mammals, taxonomic turnover was responsible for most of these dissimilarities, providing strong evidence of an estuarine gradient, as revealed by the taxonomic composition detected by eDNA metabarcoding. Beta diversity analyses also showed a significant separation of station 5 (freshwater) from the others, particularly in terms of phylogenetic and functional diversity. Stations 2 to 4 showed a tighter clustering, while station 1 appeared to be intermediate. However, for birds, the most peripheral station was actually station 1. This could be explained by the low number of bird detections, as species more closely associated with terrestrial environments were absent in station 1. Additionally, a stronger dilution effect in the marine environment (Taberlet et al., 2018) may have contributed to this pattern. Teichert et al. (2018a) studied actinopterygian communities in French estuaries of the North East Atlantic and explored the influence of various environmental factors on the taxonomic, functional, and phylogenetic diversity of actinopterygian assemblages. Their main conclusions suggested that actinopterygian diversity was primarily driven by shifts in species composition along estuarine gradients, whereas functional dissimilarity was constrained both spatially and temporally. This pattern highlighted significant changes in taxonomic composition along estuarine gradients, while spatial functional dissimilarity remained limited, which may result from the dominance of few species sharing similar ecological traits along estuaries (Villéger et al., 2013).

There was no clear separation in the taxa detected on either side of the TPP dam in this study, except for birds. In this case, the patterns appeared to be linked to detection capacities influenced by habitat and species-specific ecological traits. The results did not indicate any TPP dam effect on species diversity when comparing the actinopterygian and mammal taxa detected on each side of the dam. In fact, stations 1 and 2 shared 41 MOTUs, including four marine mammal taxa (Phocidae, two Delphininae, and *Phocoena phocoena*) as well as at least 13 actinopterygian species ecologically associated with the marine environment. While the results of this study are not sufficient to definitively confirm connectivity across the TPP dam, they are consistent with previous results by Rault et al. (2023), which demonstrated ecological connectivity across the dam for the ichthyoplankton assemblage.

Upstream, at the “Le Châtelier” lock, there is conversely clear evidence of a distinct biodiversity pattern between each side of the structure, with station 5 consistently exhibiting the most divergent alpha diversity values. Similarly, beta diversity analyses for actinopterygians and mammals revealed clear separation between station 5 and the others. This difference is primarily due to station 5 being a strictly freshwater site, but the effect is likely amplified by the Le Châtelier lock, which acts as a barrier to species movement and creates a salinity gradient between the upstream and downstream sections.

### 3. eDNA Metabarcoding as a Tool for Estuarine Biomonitoring

Assessing the ecological health of aquatic ecosystems is crucial in the current context of biodiversity loss to guide and prioritize management actions (e.g. Madon et al., 2023; Teichert et al., 2018b). In our study, species occurrences detected through eDNA metabarcoding allowed us to compute a set of non quantitative metrics of biodiversity such as species richness (SR), functional richness (FRic), standardized effect size of functional richness (SES.FRic), phylogenetic diversity (PD), and standardized effect size of phylogenetic diversity (SES.PD).

These metrics complement traditional tools used to assess estuary quality, such as the ELFI index developed for the European Water Framework Directive (WFD), which already incorporates a functional component with species traits. However, dedicated functional diversity indices can provide additional insights into the effects of local disturbances on actinopterygian assemblages. For instance, Teichert et al. (2018b) showed that functional redundancy tends to decrease in impacted estuaries and may be more sensitive to anthropogenic pressures than species richness or abundance alone. Integrating taxonomic, phylogenetic, and functional approaches offers valuable insights into the processes of structuring biological assemblages. The complementarity of these approaches has broad applications in macroecology and can enhance our understanding of ecosystem dynamics (Cardoso et al., 2014; Villéger et al., 2013).

The species lists generated based on the WFD, AnaCoNoR, and eDNA metabarcoding (this study) surveys, exhibit substantial differences due to variations in the protocols applied, techniques used (which do not allow sampling in the exact same stations of the estuary), the sampling effort, the time of year, and the species identification methods. Despite these methodological differences, no significant differences were observed in diversity metrics between the three surveys. In particular, eDNA metabarcoding produced consistent values that were not markedly different from those obtained in WFD and AnaCoNoR studies. The only notable difference in metric calculations was observed for AnaCoNoR, where standardized effect size of phylogenetic diversity (SES.PD_actino) was significantly lower than expected given the number of detected taxa. These results are consistent with the AnaCoNoR survey, which documented a decrease in marine and diadromous juvenile detections in the Rance (Rault et al., 2023).

eDNA metabarcoding is particularly well suited for assessing phylogenetic diversity (Diniz-Filho et al., 2024), as it generates comprehensive taxonomic inventories, including rare and cryptic taxa, and does not rely on quantitative abundance data or life stage identification. These limitations, especially the lack of trait information by life stage, are more restrictive for functional diversity assessments. In our case, this was partly addressed using a trait database tailored to Northeast Atlantic estuarine species (Teichert et al., 2018a). Given our focus on local spatio-temporal comparisons, we used diversity metrics across taxonomic, phylogenetic, and functional dimensions. To serve as biodiversity indicators, however, these metrics must be interpreted within a reference framework and tied to clear management objectives (Arpin et al., 2025).

Further sampling campaigns and analyses are needed to confirm these conclusions, but these initial results suggest that eDNA metabarcoding can serve as a suitable complementary tool for monitoring mobile fauna biodiversity in the Rance estuary. The sampling protocol of this approach is less constrained by depth parameters, is non-invasive and non-destructive, and provides a comprehensive snapshot of the vertebrate community, encompassing actinopterygians, marine and terrestrial mammals, amphibians, and birds. Additionally, this study proposes a set of diversity metrics that capture different aspects of taxonomic, phylogenetic, and functional diversity, derived from eDNA metabarcoding species lists, offering a more integrative perspective on estuarine biodiversity.

## Acknowledgements

We thank the technical staff of the Marine Station of Dinard (Muséum national d’Histoire naturelle) for their support during fieldwork and assistance with sampling. We are also grateful to the SPYGEN laboratory, and especially to Alice Valentini, for their work and the valuable discussions regarding the methodology.

## Data Availability Statement

Occurrence data are available online via GBIF at [https://www.gbif.org/dataset/787386a7-aa69-4f99-ba95-9fd5a700705c], and the code used for the analyses is accessible on GitHub at [https://github.com/HaderleRachel/eDNA_Rance_river].

## Ethics Approval Statement

The authors confirm that the manuscript has not been submitted elsewhere, and that all research meets the ethical guidelines of the study country.

## References

Ahn, H., Kume, M., Terashima, Y., Ye, F., Kameyama, S., Miya, M., Yamashita, Y., Kasai, A., 2020. Evaluation of fish biodiversity in estuaries using environmental DNA metabarcoding. PLoS One 15, e0231127.

Albouy, C., Leprieur, F., Le Loc’h, F., Mouquet, N., Meynard, C.N., Douzery, E.J.P., Mouillot, D., 2015. Projected impacts of climate warming on the functional and phylogenetic components of coastal Mediterranean fish biodiversity. Ecography 38, 681–689. 10.1111/ecog.01254

Andersson, A., Bissett, A., Finstad, A., Fossøy, F., Grosjean, M., Hope, M., Jeppesen, T., Kõljalg, U., Lundin, D., Nilsson, R., Prager, M., Svenningsen, C., Schigel, D., 2021. Publishing DNA-derived data through biodiversity data platforms. v1.0 Copenhagen: GBIF Secretariat. Copenhagen: GBIF Secretariat. 10.35035/doc-vf1a-nr22

Arevalo, E., Cabral, H.N., Villeneuve, B., Possémé, C., Lepage, M., 2023. Fish larvae dynamics in temperate estuaries: A review on processes, patterns and factors that determine recruitment. Fish and Fisheries 24, 466–487. 10.1111/faf.12740

Arpin, I., Kurek, M., Therville, C., Paillet, Y., 2025. Comprendre la fabrique de l’utilisabilité des indicateurs de biodiversité. Le cas des dendromicrohabitats. Naturae 2025. 10.5852/naturae2025a8

Attrill, M.J., 2002. A testable linear model for diversity trends in estuaries. Journal of Animal Ecology 71, 262–269. 10.1046/j.1365-2656.2002.00593.x

Baselga, A., 2012. The relationship between species replacement, dissimilarity derived from nestedness, and nestedness. Global Ecology and Biogeography 21, 1223–1232. 10.1111/j.1466-8238.2011.00756.x

Birk, S., Bonne, W., Borja, A., Brucet, S., Courrat, A., Poikane, S., Solimini, A., van de Bund, W., Zampoukas, N., Hering, D., 2012b. Three hundred ways to assess Europe’s surface waters: An almost complete overview of biological methods to implement the Water Framework Directive. Ecological Indicators 18, 31–41. 10.1016/j.ecolind.2011.10.009

Bohmann, K., Evans, A., Gilbert, M.T.P., Carvalho, G.R., Creer, S., Knapp, M., Yu, D.W., de Bruyn, M., 2014. Environmental DNA for wildlife biology and biodiversity monitoring. Trends in Ecology & Evolution 29, 358– 367. 10.1016/j.tree.2014.04.003

Boyer, F., Mercier, C., Bonin, A., Le Bras, Y., Taberlet, P., Coissac, E., 2016. obitools: a unix-inspired software package for DNA metabarcoding. Mol Ecol Resour 16, 176–182. 10.1111/1755-0998.12428

Bylemans, J., Gleeson, D.M., Duncan, R.P., Hardy, C.M., Furlan, E.M., 2019. A performance evaluation of targeted eDNA and eDNA metabarcoding analyses for freshwater fishes. Environmental DNA 1, 402–414. 10.1002/edn3.41

Cardoso, P., Rigal, F., Borges, P.A.V., Carvalho, J.C., 2014. A new frontier in biodiversity inventory: a proposal for estimators of phylogenetic and functional diversity. Methods in Ecology and Evolution 5, 452–461. 10.1111/2041-210X.12173

Carpentier, A., Feunteun, E., Acou, A., 2014. Directive cadre eau - suivi ichtyologique des masses d’eau de transition : compte rendu des opérations de pêche sur l’estuaire.

Carpentier, A., Feunteun, E., Acou, A., 2013. Directive cadre eau - suivi ichtyologique des masses d’eau de transition : compte rendu des opérations de pêche sur l’estuaire.

Carpentier, A., Feunteun, E., Acou, A., 2012. Directive cadre eau - suivi ichtyologique des masses d’eau de transition : compte rendu des opérations de pêche sur l’estuaire.

Chiquillo, K.L., Wong, J.M., Eirin-Lopez, J.M., 2024. Ecological forensic testing: Using multiple primers for eDNA detection of marine vertebrates in an estuarine lagoon subject to anthropogenic influences. Gene 928, 148720. 10.1016/j.gene.2024.148720

Coates, S., Waugh, A., Anwar, A., Robson, M., 2007. Efficacy of a multi-metric fish index as an analysis tool for the transitional fish component of the Water Framework Directive. Marine Pollution Bulletin, Implementation of the Water Framework Directive in European marine waters 55, 225–240. 10.1016/j.marpolbul.2006.08.029

Cole, V.J., Harasti, D., Lines, R., Stat, M., 2022. Estuarine fishes associated with intertidal oyster reefs characterized using environmental DNA and baited remote underwater video. Environmental DNA 4, 50–62. 10.1002/edn3.190

Costanza, R., Kemp, W.M., Boynton, W.R., 1993. Predictability, scale, and biodiversity in coastal and estuarine ecosystems: implications for management. Ambio 88–96.

Deiner, K., Altermatt, F., 2014. Transport Distance of Invertebrate Environmental DNA in a Natural River. PLOS ONE 9, e88786. 10.1371/journal.pone.0088786

Deiner, K., Fronhofer, E.A., Mächler, E., Walser, J.-C., Altermatt, F., 2016. Environmental DNA reveals that rivers are conveyer belts of biodiversity information. Nat Commun 7, 12544. 10.1038/ncomms12544

Delpech, C., Courrat, A., Pasquaud, S., Lobry, J., Le Pape, O., Nicolas, D., Boët, P., Girardin, M., Lepage, M., 2010. Development of a fish-based index to assess the ecological quality of transitional waters: The case of French estuaries. Marine Pollution Bulletin 60, 908–918. 10.1016/j.marpolbul.2010.01.001

Diniz-Filho, J.A.F., Bini, L.M., Targueta, C.P., Telles, M.P. de C., Jardim, L., Machado, K.B., Nabout, J.C., Nunes, R., Vieira, L.C.G., Soares, T.N., 2024. Environmental DNA and biodiversity patterns: a call for a community phylogenetics approach. Perspectives in Ecology and Conservation 22, 15–23. 10.1016/j.pecon.2024.01.006

Dray, S., Blanchet, G., Borcard, D., Guenard, G., Jombart, T., Larocque, G., Legendre, P., Madi, N., Wagner, H.H., Dray, M.S., 2018. Package ‘adespatial.’ R package 2018, 3–8.

Dray, S., Pélissier, R., Couteron, P., Fortin, M.-J., Legendre, P., Peres-Neto, P.R., Bellier, E., Bivand, R., Blanchet, F.G., De Cáceres, M., Dufour, A.-B., Heegaard, E., Jombart, T., Munoz, F., Oksanen, J., Thioulouse, J., Wagner, H.H., 2012. Community ecology in the age of multivariate multiscale spatial analysis. Ecological Monographs 82, 257–275. 10.1890/11-1183.1

Elliott, M., Quintino, V., 2007. The Estuarine Quality Paradox, Environmental Homeostasis and the difficulty of detecting anthropogenic stress in naturally stressed areas. Marine Pollution Bulletin 54, 640–645. 10.1016/j.marpolbul.2007.02.003

Elliott, M., Whitfield, A.K., Potter, I.C., Blaber, S.J.M., Cyrus, D.P., Nordlie, F.G., Harrison, T.D., 2007. The guild approach to categorizing estuarine fish assemblages: a global review. Fish and Fisheries 8, 241–268. 10.1111/j.1467-2679.2007.00253.x

Euclide, P.T., Lor, Y., Spear, M.J., Tajjioui, T., Vander Zanden, J., Larson, W.A., Amberg, J.J., 2021. Environmental DNA metabarcoding as a tool for biodiversity assessment and monitoring: reconstructing established fish communities of north-temperate lakes and rivers. Diversity and Distributions 27, 1966–1980. 10.1111/ddi.13253

Faith, D.P., 1992. Conservation evaluation and phylogenetic diversity. Biological Conservation 61, 1–10. 10.1016/0006-3207(92)91201-3

Faith, D.P., Baker, A.M., 2006. Phylogenetic Diversity (PD) and Biodiversity Conservation: Some Bioinformatics Challenges. Evol Bioinform Online 2, 117693430600200007. 10.1177/117693430600200007

García-Machado, E., Laporte, M., Normandeau, E., Hernández, C., Côté, G., Paradis, Y., Mingelbier, M., Bernatchez, L., 2022. Fish community shifts along a strong fluvial environmental gradient revealed by eDNA metabarcoding. Environmental DNA 4, 117–134. 10.1002/edn3.221

GBIF Secretariat, 2024. Metabarcoding Data Toolkit – user guide. Copenhagen: Global Biodiversity Facility. 10.35035/doc-wkpc-m352

Gibson, T.I., Carvalho, G., Ellison, A., Gargiulo, E., Hatton-Ellis, T., Lawson-Handley, L., Mariani, S., Collins, R.A., Sellers, G., Antonio Distaso, M., Zampieri, C., Creer, S., 2023. Environmental DNA metabarcoding for fish diversity assessment in a macrotidal estuary: A comparison with established fish survey methods. Estuarine, Coastal and Shelf Science 294, 108522. 10.1016/j.ecss.2023.108522

Gotelli, N.J., McCabe, D.J., 2002. Species Co-Occurrence: A Meta-Analysis of J. M. Diamond’s Assembly Rules Model. Ecology 83, 2091–2096. 10.1890/0012-9658(2002)083[2091:SCOAMA]2.0.CO;2

Günther, B., Jourdain, E., Rubincam, L., Karoliussen, R., Cox, S.L., Arnaud Haond, S., 2022. Feces DNA analyses track the rehabilitation of a free-ranging beluga whale. Sci Rep 12, 6412. 10.1038/s41598-022-09285-8

Haderlé, R., Bouveret, L., Chazal, J., Girardet, J., Iglésias, S., Lopez, P.-J., Millon, C., Valentini, A., Ung, V., Jung, J.-L., 2024a. eDNA-based survey of the marine vertebrate biodiversity off the west coast of Guadeloupe (French West Indies). Biodiversity Data Journal 12, e125348. 10.3897/BDJ.12.e125348

Haderlé, R., Ung, V., Jung, J.-L., 2024b. VeTAPRH: A Taxonomic Assignment Protocol for Vertebrates Applied to eDNA Metabarcoding Data, Including Molecular, Taxonomic and Ecological Criteria. Biodiversity Information Science and Standards 8. 10.3897/biss.8.141746

Harper, L.R., Handley, L., Carpenter, A., Ghazali, M., Muri, C., Macgregor, C.J., Logan, T.W., Law, A., Breithaupt, T., Read, D., Mcdevitt, A., Hänfling, B., 2019. Environmental DNA (eDNA) metabarcoding of pond water as a tool to survey conservation and management priority mammals. bioRxiv null, null. 10.1101/546218

Harrison, T.D., Kelly, F.L., 2013. Development of an estuarine multi-metric fish index and its application to Irish transitional waters. Ecological Indicators 34, 494–506. 10.1016/j.ecolind.2013.06.018

Henriques, S., Guilhaumon, F., Villéger, S., Amoroso, S., França, S., Pasquaud, S., Cabral, H.N., Vasconcelos, R.P., 2017. Biogeographical region and environmental conditions drive functional traits of estuarine fish assemblages worldwide. Fish and Fisheries 18, 752–771. 10.1111/faf.12203

Inventaire National du Patrimoine Naturel, 2023. Occurrence datasetgic [WWW Document]. URL https://www.gbif.org/occurrence/3844758827 (accessed 7.11.25).

Ip, J.C.-H., Loke, H.-X., Yiu, S.K.F., Zhao, M., Li, Y., Lin, Y., How, C.-M., Mo, J., Yan, M., Cheng, J., Lai, V.C.-S., Chan, L.L., Leung, K.M.Y., Qiu, J.-W., 2024. Bottom Trawling and Multi-Marker eDNA Metabarcoding Surveys Reveal Highly Diverse Vertebrate and Crustacean Communities: A Case Study in an Urbanized Subtropical Estuary. Environmental DNA 6, e70031. 10.1002/edn3.70031

Jackman, J.M., Benvenuto, C., Coscia, I., Carvalho, C.O., Ready, J., Boubli, J., Magnusson, W., Mcdevitt, A., Sales, N., 2021. eDNA in a bottleneck: obstacles to fish metabarcoding studies in megadiverse freshwater systems. bioRxiv null, null. 10.1101/2021.01.05.425493

Jane, S.F., Wilcox, T.M., McKelvey, K.S., Young, M.K., Schwartz, M.K., Lowe, W.H., Letcher, B.H., Whiteley, A.R., 2015. Distance, flow and PCR inhibition: e DNA dynamics in two headwater streams. Molecular Ecology Resources 15, 216–227. 10.1111/1755-0998.12285

Jetz, W., Thomas, G.H., Joy, J.B., Hartmann, K., Mooers, A.O., 2012. The global diversity of birds in space and time. Nature 491, 444–448. 10.1038/nature11631

Jin, Y., Qian, H., 2023. U. PhyloMaker: An R package that can generate large phylogenetic trees for plants and animals. Plant Diversity 45, 347–352.

Jung, J.-L., 2024. Environmental DNA for observing marine mammals in the marine protected areas of Iroise and the Antilles, in: L’inventaire de La Biodiversité, V. Colin. Paris.

Jung, J.-L., Gicquel, C., 2021. Projet CetADNe : Détection de mammifères marins par analyse d’ADN environnemental dans des prélèvements d’eau de mer. Résultats de la première campagne test en mer d’Iroise (report). UMR ISYEB - MNHN.

Kassambara, A., Mundt, F., 2017. Package ‘factoextra.’ Extract and visualize the results of multivariate data analyses 76, 10–18637.

Kembel, S.W., Ackerly, D.D., Blomberg, S.P., Cornwell, W.K., Cowan, P.D., Helmus, M.R., Morlon, H., Webb, C.O., 2014. Package ‘picante.’ R Foundation for Statistical Computing, Vienna, Austria: https://cran.r-project.

Kennish, M.J., 2002. Environmental threats and environmental future of estuaries. Environmental conservation 29, 78–107.

Laporte, M., Reny-Nolin, E., Chouinard, V., Hernandez, C., Normandeau, É., Bougas, B., Côté, C., Behmel, S., Bernatchez, L., 2021. Proper environmental DNA metabarcoding data transformation reveals temporal stability of fish communities in a dendritic river system. Environmental DNA null, null. 10.1002/edn3.224

Le Mao, P., 1986. Feeding relationships between the benthic infauna and the dominant benthic fish of the Rance Estuary (France). Journal of the Marine Biological Association of the United Kingdom 66, 391–401.

Leprieur, F., Albouy, C., Bortoli, J.D., Cowman, P.F., Bellwood, D.R., Mouillot, D., 2012. Quantifying Phylogenetic Beta Diversity: Distinguishing between ‘True’ Turnover of Lineages and Phylogenetic Diversity Gradients. PLOS ONE 7, e42760. 10.1371/journal.pone.0042760

Mächler, E., Little, C.J., Wüthrich, R., Alther, R., Fronhofer, E.A., Gounand, I., Harvey, E., Hürlemann, S., Walser, J., Altermatt, F., 2019. Assessing different components of diversity across a river network using eDNA. Environmental DNA 1, 290–301. 10.1002/edn3.33

Madon, B., David, R., Torralba, A., Jung, A., Marengo, M., Thomas, H., 2023. A review of biodiversity research in ports: Let’s not overlook everyday nature! Ocean & Coastal Management 242, 106623. 10.1016/j.ocecoaman.2023.106623

Madon, B., Haderlé, R., Emma, A., Romain, D., Quentin, F., Michel, M., Hélène, T., Antonio, T., Alice, V., Jung, J.-L., In Press. eDNA and Citizen Science Reveal Hidden Fish Biodiversity in Climate-Stressed Urban Ports of the Mediterranean Sea. Environmental DNA. doi: 10.1002/edn3.70142

Magneville, C., Loiseau, N., Albouy, C., Casajus, N., Claverie, T., Escalas, A., Leprieur, F., Maire, E., Mouillot, D., Villéger, S., 2022. mFD: an R package to compute and illustrate the multiple facets of functional diversity. Ecography 2022. 10.1111/ecog.05904

Mazel, F., Pennell, M.W., Cadotte, M.W., Diaz, S., Dalla Riva, G.V., Grenyer, R., Leprieur, F., Mooers, A.O., Mouillot, D., Tucker, C.M., 2018. Prioritizing phylogenetic diversity captures functional diversity unreliably. Nature communications 9, 2888.

McCormick, S.D., Hansen, L.P., Quinn, T.P., Saunders, R.L., 1998. Movement, migration, and smolting of Atlantic salmon (Salmo salar). Can. J. Fish. Aquat. Sci. 55, 77–92. 10.1139/d98-011

McLusky, D.S., Elliott, M., 2004. The estuarine ecosystem: ecology, threats and management. OUP Oxford.

Pavkovic, M., Carpentier, A., Duhamel, S., Carassou, L., Lobry, J., Feunteun, E., Teichert, N., 2025. Estuarine lateral ecotones shape taxonomic and functional structure of fish assemblages. The case of the Seine Estuary, France. Estuarine, Coastal and Shelf Science 313, 109066. 10.1016/j.ecss.2024.109066

Pérez-Domínguez, R., Maci, S., Courrat, A., Lepage, M., Borja, A., Uriarte, A., Neto, J.M., Cabral, H., St.Raykov, V., Franco, A., Alvarez, M.C., Elliott, M., 2012. Current developments on fish-based indices to assess ecological-quality status of estuaries and lagoons. Ecological Indicators 23, 34–45. 10.1016/j.ecolind.2012.03.006

Polanco, A.P., Martinezguerra, M.M., Marques, V., Villa-Navarro, F., Pérez, G.H.B., Cheutin, M.-C., Dejean, T., Hocdé, R., Juhel, J., Maire, E., Manel, S., Spescha, M., Valentini, A., Mouillot, D., Albouy, C., Pellissier, L., 2021. Detecting aquatic and terrestrial biodiversity in a tropical estuary using environmental DNA. Biotropica 53, 1606–1619. 10.1111/btp.13009

Potter, I.C., Tweedley, J.R., Elliott, M., Whitfield, A.K., 2015. The ways in which fish use estuaries: a refinement and expansion of the guild approach. Fish and Fisheries 16, 230–239. 10.1111/faf.12050

R Core Team, 2025. R: A Language and Environment for Statistical Computing. R Foundation for Statistical Computing, Vienna, Austria. [WWW Document]. URL https://www.r-project.org/ (accessed 1.23.25).

Rabosky, D.L., Chang, J., Cowman, P.F., Sallan, L., Friedman, M., Kaschner, K., Garilao, C., Near, T.J., Coll, M., Alfaro, M.E., 2018. An inverse latitudinal gradient in speciation rate for marine fishes. Nature 559, 392–395.

Rault, P., Teichert, N., Feunteun, E., Desroy, N., Lepage, M., Kervarec, G., Chapalain, M., Husset, M.-C., Prod’homme, J., Trancart, T., Acou, A., Carpentier, A., 2023. Projet d’étude AnaCoNoR : Analyse de la Connectivité et de la fonction de Nourricerie pour les jeunes stades de poissons du bassin de la Rance. 10.13140/RG.2.2.31096.21761

Retiere, C., 1994. Tidal power and the aquatic environment of La Rance. Biological Journal of the Linnean Society 51, 25–36.

Rey, A., Viard, F., Lizé, A., Corre, E., Valentini, A., Thiriet, P., 2023. Coastal rocky reef fish monitoring in the context of the Marine Strategy Framework Directive: Environmental DNA metabarcoding complements underwater visual census. Ocean & Coastal Management 241, 106625. 10.1016/j.ocecoaman.2023.106625

Ruppert, K.M., Kline, R.J., Rahman, M.S., 2019. Past, present, and future perspectives of environmental DNA (eDNA) metabarcoding: A systematic review in methods, monitoring, and applications of global eDNA. Global Ecology and Conservation 17, e00547. 10.1016/j.gecco.2019.e00547

Saenz-Agudelo, P., Delrieu-Trottin, E., DiBattista, J.D., Martínez-Rincon, D., Morales-González, S., Pontigo, F., Ramírez, P., Silva, A., Soto, M., Correa, C., 2022. Monitoring vertebrate biodiversity of a protected coastal wetland using eDNA metabarcoding. Environmental DNA 4, 77–92. 10.1002/edn3.200

Safi, K., Cianciaruso, M.V., Loyola, R.D., Brito, D., Armour-Marshall, K., Diniz-Filho, J.A.F., 2011. Understanding global patterns of mammalian functional and phylogenetic diversity. Phil. Trans. R. Soc. B 366, 2536–2544. 10.1098/rstb.2011.0024

Stoeckle, M.Y., Soboleva, L., Charlop-Powers, Z., 2017. Aquatic environmental DNA detects seasonal fish abundance and habitat preference in an urban estuary. PLOS ONE 12, e0175186. 10.1371/journal.pone.0175186

Taberlet, P., Bonin, A., Zinger, L., Coissac, É., 2018. Environmental DNA: For Biodiversity Research and Monitoring, Environmental DNA: For Biodiversity Research and Monitoring. 10.1093/oso/9780198767220.001.0001

Takahashi, M., Frøslev, T.G., Paupério, J., Thalinger, B., Klymus, K., Helbing, C.C., Villacorta-Rath, C., Silliman, K., Thompson, L.R., Jungbluth, S.P., Yong, S.Y., Formel, S., Jenkins, G., Laporte, M., Deagle, B., Rajbhandari, S., Jeppesen, T.S., Bissett, A., Jerde, C., Hahn, E.E., Schriml, L.M., Hunter, C., Newman, P., Woollard, P., Harper, L.R., Dunn, N., West, K., Haderlé, R., Wilkinson, S., Acharya-Patel, N., Lopez, M.L.D., Cochrane, G., Berry, O., 2025. A Metadata Checklist and Data Formatting Guidelines to Make eDNA FAIR (Findable, Accessible, Interoperable, and Reusable). Environmental DNA 7, e70100. 10.1002/edn3.70100

Teichert, N., Lepage, M., Chevillot, X., Lobry, J., 2018a. Environmental drivers of taxonomic, functional and phylogenetic diversity (alpha, beta and gamma components) in estuarine fish communities. Journal of Biogeography 45, 406–417. 10.1111/jbi.13133

Teichert, N., Lepage, M., Lobry, J., 2018c. Beyond classic ecological assessment: The use of functional indices to indicate fish assemblages sensitivity to human disturbance in estuaries. Science of The Total Environment 639, 465–475. 10.1016/j.scitotenv.2018.05.179

Trancart, T., Teichert, N., Lamoureux, J., Gharnit, E., Acou, A., De Oliveira, E., Roy, R., Feunteun, E., 2022b. A possible strong impact of tidal power plant on silver eels’ migration. Estuarine, Coastal and Shelf Science 278, 108116. 10.1016/j.ecss.2022.108116

Upham, N.S., Esselstyn, J.A., Jetz, W., 2019. Inferring the mammal tree: Species-level sets of phylogenies for questions in ecology, evolution, and conservation. PLOS Biology 17, e3000494. 10.1371/journal.pbio.3000494

Valentini, A., Pompanon, F., Taberlet, P., 2009. DNA barcoding for ecologists. Trends in Ecology & Evolution 24, 110–117. 10.1016/j.tree.2008.09.011

Valentini, A., Taberlet, P., Miaud, C., Civade, R., Herder, J., Thomsen, P.F., Bellemain, E., Besnard, A., Coissac, E., Boyer, F., Gaboriaud, C., Jean, P., Poulet, N., Roset, N., Copp, G.H., Geniez, P., Pont, D., Argillier, C., Baudoin, J.-M., Peroux, T., Crivelli, A.J., Olivier, A., Acqueberge, M., Le Brun, M., Møller, P.R., Willerslev, E., Dejean, T., 2016. Next-generation monitoring of aquatic biodiversity using environmental DNA metabarcoding. Molecular Ecology 25, 929–942. 10.1111/mec.13428

Villéger, S., Grenouillet, G., Brosse, S., 2013. Decomposing functional β-diversity reveals that low functional β-diversity is driven by low functional turnover in European fish assemblages. Global Ecology and Biogeography 22, 671–681. 10.1111/geb.12021

Villéger, S., Mason, N.W.H., Mouillot, D., 2008. New Multidimensional Functional Diversity Indices for a Multifaceted Framework in Functional Ecology. Ecology 89, 2290–2301. 10.1890/07-1206.1

Whitfield, A.K., 2015. Why are there so few freshwater fish species in most estuaries? Journal of Fish Biology 86, 1227–1250. 10.1111/jfb.12641

Whitfield, A.K., Elliott, M., Basset, A., Blaber, S.J.M., West, R.J., 2012. Paradigms in estuarine ecology–a review of the Remane diagram with a suggested revised model for estuaries. Estuarine, Coastal and Shelf Science 97, 78–90.

Wieczorek, J., Bloom, D., Guralnick, R., Blum, S., Döring, M., Giovanni, R., Robertson, T., Vieglais, D., 2012. Darwin Core: An Evolving Community-Developed Biodiversity Data Standard. PLOS ONE 7, e29715. 10.1371/journal.pone.0029715

Wiens, J.J., Ackerly, D.D., Allen, A.P., Anacker, B.L., Buckley, L.B., Cornell, H.V., Damschen, E.I., Jonathan Davies, T., Grytnes, J., Harrison, S.P., Hawkins, B.A., Holt, R.D., McCain, C.M., Stephens, P.R., 2010. Niche conservatism as an emerging principle in ecology and conservation biology. Ecology Letters 13, 1310–1324. 10.1111/j.1461-0248.2010.01515.x

Wiens, J.J., Graham, C.H., 2005. Niche Conservatism: Integrating Evolution, Ecology, and Conservation Biology. Annu. Rev. Ecol. Evol. Syst. 36, 519–539. 10.1146/annurev.ecolsys.36.102803.095431

Wilman, H., Belmaker, J., Simpson, J., de la Rosa, C., Rivadeneira, M.M., Jetz, W., 2014. EltonTraits 1.0: Species-level foraging attributes of the world’s birds and mammals. Ecology 95, 2027–2027. 10.1890/13-1917.1

Yamanaka, H., Minamoto, T., 2016. The use of environmental DNA of fishes as an efficient method of determining habitat connectivity. Ecological Indicators 62, 147–153. 10.1016/j.ecolind.2015.11.022

Yilmaz, P., Kottmann, R., Field, D., Knight, R., Cole, J.R., Amaral-Zettler, L., Gilbert, J.A., Karsch-Mizrachi, I., Johnston, A., Cochrane, G., Vaughan, R., Hunter, C., Park, J., Morrison, N., Rocca-Serra, P., Sterk, P., Arumugam, M., Bailey, M., Baumgartner, L., Birren, B.W., Blaser, M.J., Bonazzi, V., Booth, T., Bork, P., Bushman, F.D., Buttigieg, P.L., Chain, P.S.G., Charlson, E., Costello, E.K., Huot-Creasy, H., Dawyndt, P., DeSantis, T., Fierer, N., Fuhrman, J.A., Gallery, R.E., Gevers, D., Gibbs, R.A., Gil, I.S., Gonzalez, A., Gordon, J.I., Guralnick, R., Hankeln, W., Highlander, S., Hugenholtz, P., Jansson, J., Kau, A.L., Kelley, S.T., Kennedy, J., Knights, D., Koren, O., Kuczynski, J., Kyrpides, N., Larsen, R., Lauber, C.L., Legg, T., Ley, R.E., Lozupone, C.A., Ludwig, W., Lyons, D., Maguire, E., Methé, B.A., Meyer, F., Muegge, B., Nakielny, S., Nelson, K.E., Nemergut, D., Neufeld, J.D., Newbold, L.K., Oliver, A.E., Pace, N.R., Palanisamy, G., Peplies, J., Petrosino, J., Proctor, L., Pruesse, E., Quast, C., Raes, J., Ratnasingham, S., Ravel, J., Relman, D.A., Assunta-Sansone, S., Schloss, P.D., Schriml, L., Sinha, R., Smith, M.I., Sodergren, E., Spor, A., Stombaugh, J., Tiedje, J.M., Ward, D.V., Weinstock, G.M., Wendel, D., White, O., Whiteley, A., Wilke, A., Wortman, J.R., Yatsunenko, T., Glöckner, F.O., 2011. Minimum information about a marker gene sequence (MIMARKS) and minimum information about any (x) sequence (MIxS) specifications. Nat Biotechnol 29, 415–420. 10.1038/nbt.1823

Yong, S.Y., Takahashi, M., 2025. FAIRe-fier: FAIR eDNA metadata verifier. v2. CSIRO. Service Collection.

Zainal Abidin, D.H., Mohd. Nor, S.A., Lavoué, S., A. Rahim, M., Mohammed Akib, N.A., 2022. Assessing a megadiverse but poorly known community of fishes in a tropical mangrove estuary through environmental DNA (eDNA) metabarcoding. Scientific Reports 12, 16346.

Zhang, S., Zhao, J., Yao, M., 2023b. Urban landscape-level biodiversity assessments of aquatic and terrestrial vertebrates by environmental DNA metabarcoding. Journal of Environmental Management 340, 117971. 10.1016/j.jenvman.2023.117971

