## Appendix A for "A Multi-Taxa Approach to Estuarine Biomonitoring: Assessing Vertebrate Biodiversity and Ecological Continuity using Environmental DNA Metabarcoding in the Rance River (Brittany, France)"

*__Appendix A__ List of hypothetical species associated with each identified MOTU.*

| Class | Identified taxon | Possible taxa (a: absent from reference databases; b: similar DNA but taxon not locally known) | STATION 1 | STATION 2 | STATION 3 | STATION 4 | STATION 5 |
|---|---|---|---|---|---|---|---|
| Actinopterygii | Perciformes | Pholis gunnellus, Taurulus bubalis | 1 | 0 | 0 | 0 | 0 |
| Actinopterygii | Anguilla_anguilla_1 | | 1 | 0 | 1 | 0 | 0 |
| Actinopterygii | Anguilla_anguilla_2 | | 0 | 1 | 1 | 1 | 1 |
| Actinopterygii | Atherina_presbyter | | 1 | 1 | 1 | 1 | 0 |
| Actinopterygii | Belone_belone | | 1 | 1 | 0 | 1 | 0 |
| Actinopterygii | Coryphoblennius_galerita | | 1 | 0 | 0 | 0 | 0 |
| Actinopterygii | Lipophrys_pholis | | 1 | 1 | 1 | 0 | 0 |
| Actinopterygii | Parablennius_gattorugine | | 1 | 1 | 1 | 1 | 0 |
| Actinopterygii | Callionymus_lyra | | 1 | 1 | 1 | 1 | 0 |
| Actinopterygii | Lepomis_gibbosus | | 0 | 1 | 0 | 0 | 1 |
| Actinopterygii | Sardina_pilchardus | | 1 | 1 | 1 | 1 | 0 |
| Actinopterygii | Clupeidae | Clupea harengus, Sprattus sprattus | 1 | 1 | 1 | 1 | 0 |
| Actinopterygii | Cyprinidae | Gobio gobio, Alburnus alburnus, Alburnoides bipunctatus, Leuciscus leuciscus | 0 | 1 | 1 | 1 | 1 |
| Actinopterygii | Cyprinidae_1 | Leuciscus leuciscus, Leuciscus burdigalensis, Scardinius erythrophtalmus, Rutilus rutilus, Alburnus alburnus, Alburnoides bipunctatus, Tinca tinca | 0 | 1 | 0 | 0 | 1 |
| Actinopterygii | Cyprinidae_2 | Tinca tinca, Ctenopharyngodon idella, Scardinius erythrophtalmus | 0 | 0 | 1 | 0 | 1 |
| Actinopterygii | Cyprinidae_3 | Rutilus rutilus, Alburnus alburnus, Alburnoides bipunctatus, Scardinius erythrophtalmus | 0 | 0 | 0 | 0 | 1 |
| Actinopterygii | Cyprininae | Carassius auratus, Carassius carassius, Carassius gibelio, Cyprinus carpio | 0 | 0 | 0 | 1 | 1 |
| Actinopterygii | Cyprinus_carpio | | 1 | 1 | 1 | 1 | 1 |
| Actinopterygii | Leuciscinae | Abramis brama, Blicca bjoerkna | 1 | 1 | 1 | 1 | 1 |

| | | | | | | | |
|---|---|---|---|---|---|---|---|
| Actinopterygii | Squalius_cephalus | | 0 | 0 | 1 | 1 | 1 |
| Actinopterygii | Barbatula_barbatula | | 0 | 0 | 1 | 1 | 1 |
| Actinopterygii | Esox_lucius | | 0 | 0 | 1 | 1 | 1 |
| Actinopterygii | Gaidropsaridae | Ciliata mustela, Ciliata septentrionalis (a), Enchelyopus cimbrius | 0 | 1 | 0 | 0 | 0 |
| Actinopterygii | Aphia_minuta_1 | | 1 | 1 | 0 | 0 | 0 |
| Actinopterygii | Aphia_minuta_2 | | 1 | 0 | 0 | 0 | 0 |
| Actinopterygii | Gobius_niger | | 1 | 1 | 1 | 1 | 0 |
| Actinopterygii | Gobius_paganellus_1 | | 0 | 0 | 0 | 1 | 0 |
| Actinopterygii | Gobius_paganellus_2 | | 1 | 1 | 1 | 0 | 0 |
| Actinopterygii | Pomatoschistus_sp._1 | Pomatoschistus minutus, Pomatoschistus pictus (a) | 1 | 1 | 1 | 1 | 0 |
| Actinopterygii | Pomatoschistus_sp._2 | Pomatoschistus flavescens, Pomatoschistus pictus (a) | 1 | 1 | 1 | 1 | 0 |
| Actinopterygii | Pomatoschistus_sp._3 | Pomatoschistus microps, Pomatoschistus minutus, Pomatoschistus pictus (a) | 1 | 1 | 1 | 1 | 0 |
| Actinopterygii | Chelon_sp. | Chelon labrosus, Chelon ramada, Chelon auratus | 1 | 1 | 1 | 1 | 0 |
| Actinopterygii | Mullus_surmuletus | | 1 | 1 | 0 | 0 | 0 |
| Actinopterygii | Ammodytidae_1 | Hyperoplus immaculatus, Hyperoplus lanceolatus, Ammodytes marinus, Ammodytes tobianus | 0 | 1 | 0 | 0 | 0 |
| Actinopterygii | Ammodytidae_2 | Hyperoplus immaculatus, Hyperoplus lanceolatus, Ammodytes marinus, Ammodytes tobianus | 0 | 1 | 0 | 0 | 0 |
| Actinopterygii | Ammodytidae_3 | Hyperoplus immaculatus, Hyperoplus lanceolatus, Ammodytes marinus, Ammodytes tobianus | 1 | 1 | 1 | 1 | 0 |
| Actinopterygii | Gymnammodytes_semisquamatus | | 1 | 1 | 1 | 1 | 0 |
| Actinopterygii | Centrolabrus_exoletus | | 0 | 1 | 0 | 0 | 0 |
| Actinopterygii | Ctenolabrus_rupestris | | 1 | 1 | 0 | 1 | 0 |
| Actinopterygii | Labrus_bergylta | | 1 | 1 | 1 | 1 | 0 |

| | | | | | | | |
|---|---|---|---|---|---|---|---|
| Actinopterygii | Symphodus_bailloni | | 1 | 1 | 1 | 1 | 0 |
| Actinopterygii | Symphodus_melops | | 1 | 1 | 1 | 0 | 0 |
| Actinopterygii | Dicentrarchus labrax | | 1 | 1 | 1 | 1 | 0 |
| Actinopterygii | Percidae | Perca fluviatilis, Sander lucioperca | 0 | 0 | 1 | 1 | 1 |
| Actinopterygii | Gymnocephalus_cernua | | 0 | 0 | 1 | 1 | 1 |
| Actinopterygii | Pholis_gunnellus | | 1 | 0 | 0 | 0 | 0 |
| Actinopterygii | Scomber_scombrus | | 1 | 1 | 0 | 1 | 0 |
| Actinopterygii | Diplodus_sp. | Diplodus sargus, Diplodus vulgaris (a) | 1 | 0 | 0 | 0 | 0 |
| Actinopterygii | Sparus_aurata | | 1 | 1 | 1 | 1 | 0 |
| Actinopterygii | Spondyliosoma_cantharus_1 | | 1 | 1 | 0 | 0 | 1 |
| Actinopterygii | Spondyliosoma_cantharus_2 | | 0 | 0 | 1 | 0 | 0 |
| Actinopterygii | Arnoglossus_laterna | | 1 | 0 | 0 | 0 | 0 |
| Actinopterygii | Pleuronectidae | Limanda limanda, Pleuronectes platessa, Platichthys flesus, Glyptocephalus cynoglossus, Hippoglossoides platessoides, Hippoglossus hippoglossus, Microstomus kitt | 1 | 1 | 1 | 1 | 0 |
| Actinopterygii | Zeugopterus_sp._ | Zeugopterus punctatus, Zeugopterus regius | 0 | 0 | 1 | 0 | 0 |
| Actinopterygii | Pegusa_lascaris | | 0 | 1 | 0 | 0 | 0 |
| Actinopterygii | Solea_solea | | 1 | 1 | 1 | 1 | 0 |
| Actinopterygii | Salmo trutta | Salmo trutta | 0 | 1 | 1 | 1 | 1 |
| Actinopterygii | Salmo_salar | | 1 | 1 | 1 | 1 | 0 |
| Actinopterygii | Scorpaenoidei_1 | Myoxocephalus scorpius, Taurulus bubalis, Cottunculus thomsonii, Leptagonus decagonus, Agonus cataphractus | 1 | 1 | 0 | 0 | 0 |
| Actinopterygii | Scorpaenoidei_2 | Cottus gobio, Cottus perifretum, Myoxocephalus scorpius, Cottunculus thomsonii | 0 | 0 | 0 | 1 | 1 |
| Actinopterygii | Cyclopterus_lumpus | | 1 | 1 | 0 | 0 | 0 |
| Actinopterygii | Liparis_liparis | | 1 | 0 | 0 | 0 | 0 |

| | | | | | | | |
|---|---|---|---|---|---|---|---|
| Actinopterygii | Triglinae | Eutrigla gurnardus, Chelidonichthys cuculus, Chelidonichthys lucerna, Chelidonichthys lastoviza (a), Chelidonichthys obscurus (a) | 1 | 0 | 1 | 0 | 0 |
| Actinopterygii | Hippocampus_hippocampus | | 1 | 0 | 0 | 0 | 0 |
| Actinopterygii | Nerophis_lumbriciformis | | 1 | 0 | 0 | 1 | 0 |
| Actinopterygii | Syngnathus_acus | | 0 | 0 | 1 | 0 | 0 |
| Amphibia | Bufo_spinosus | | 0 | 0 | 1 | 1 | 1 |
| Amphibia | Lissotriton_helveticus | | 0 | 1 | 0 | 0 | 1 |
| Amphibia | Salamandra_salamandra | | 0 | 0 | 0 | 0 | 1 |
| Aves | Aves | Accipiter nisus, Ardenna bulleri, Podiceps grisegena (a), Puffinus puffinus (a), Puffinus mauretanicus (a) | 0 | 0 | 1 | 0 | 1 |
| Aves | Buteo_buteo | | 0 | 0 | 0 | 0 | 1 |
| Aves | Anatidae_1 | Anser anser, Anser albifrons, Cygnus olor, Cygnus columbianus, Cygnus cygnus, Branta bernicla, Branta canadensis, Branta leucopsis (a), Branta hutchinsii (a) | 1 | 1 | 1 | 1 | 1 |
| Aves | Anatidae_2 | Anas platyrhynchos, Anas acuta, Anas crecca, Anas rubripes (a), Tadorna tadorna, Mareca strepera, Mareca penelope, Somateria mollissima, Bucephala clangula, Mergus merganser, Mergus serrator | 1 | 1 | 1 | 1 | 1 |
| Aves | Anatidae_3 | Anas platyrhynchos, Anas acuta, Anas crecca, Anas rubripes (a), Tadorna tadorna, Mareca strepera, Mareca penelope, Somateria mollissima, Bucephala clangula, Mergus merganser, Mergus serrator, Melanitta americana, Melanitta fusca (a), Melanitta nigra (a), Mergus merganser | 0 | 1 | 0 | 0 | 0 |
| Aves | Aix_galericulata | | 0 | 0 | 0 | 0 | 1 |
| Aves | Tadorna_sp. | Tadorna ferruginea, Tadorna tadorna | 0 | 0 | 1 | 1 | 0 |
| Aves | Charadriiformes | Pluvialis squatarola, Pluvialis apricaria, Pluvialis dominica, Pluvialis fulva, Stercorarius parasiticus | 0 | 0 | 1 | 0 | 0 |
| Aves | Haematopus_ostralegus_1 | | 1 | 0 | 1 | 0 | 0 |
| Aves | Haematopus_ostralegus_2 | | 0 | 1 | 0 | 0 | 0 |

| | | | | | | | |
|---|---|---|---|---|---|---|---|
| Aves | Laridae_1 | Larus argentatus, Larus marinus, Larus glaucoides, Larus canus, Larus cachinnans (a), Larus michahellis (a), Larus hyperboreus (a), Larus fuscus (a), Chroicocephalus ridibundus, Rissa tridactyla, Xema sabini, Rhodostethia rosea, Pagophila eburnea | 0 | 1 | 1 | 1 | 1 |
| Aves | Laridae_2 | Larus argentatus, Larus marinus, Larus glaucoides, Larus canus, Larus cachinnans (a), Larus michahellis (a), Larus hyperboreus (a), Larus fuscus (a), Chroicocephalus ridibundus, Rissa tridactyla, Xema sabini, Rhodostethia rosea, Pagophila eburnea | 1 | 1 | 0 | 0 | 1 |
| Aves | Scolopacidae_1 | Calidris alpina, Gallinago gallinago, Limnodromus scolopaceus | 0 | 1 | 1 | 1 | 0 |
| Aves | Scolopacidae_2 | Calidris alpina, Gallinago gallinago | 0 | 1 | 1 | 1 | 0 |
| Aves | Numenius_arquata | | 0 | 1 | 1 | 1 | 0 |
| Aves | Columbidae | Columba oenas, Columba palumbus, Streptopelia decaocto | 0 | 1 | 1 | 1 | 1 |
| Aves | Alectoris_rufa | | 0 | 0 | 0 | 1 | 0 |
| Aves | Fulica_atra | | 0 | 1 | 1 | 1 | 1 |
| Aves | Gallinula_chloropus | | 0 | 1 | 1 | 1 | 1 |
| Aves | Passeriformes_1 | Fringilla coelebs, Fringilla montifringilla, Motacilla cinerea, Motacilla tschutschensis, Motacilla alba, Piranga olivacea, Anthus hodgsoni, Anthus cervinus, Anthus godlewskii (a), Anthus gustavi (a), Anthus spinoletta (a), Anthus petrosus (a), Anthus rubescens (a), Pyrrhula pyrrhula, Carpodacus erythrinus, Pheucticus ludovicianus | 0 | 0 | 0 | 0 | 1 |
| Aves | Passeriformes_2 | Carduelis carduelis, Carduelis citrinella (a), Emberiza calandra (a), Emberiza cirlus (a), Plectrophenax nivalis | 0 | 0 | 0 | 0 | 1 |
| Aves | Passeriformes_3 | Turdus viscivorus, Turdus philomelos, Turdus iliacus, Turdus merula, Turdus pilaris (a), Parus major | 0 | 0 | 1 | 1 | 1 |

| | | | | | | | |
|---|---|---|---|---|---|---|---|
| Aves | Passeriformes_4 | Turdus philomelos, Turdus iliacus, Turdus merula, Turdus obscurus, Turdus atrogularis, Turdus ruficollis, Turdus pilaris (a), Turdus torquatus (a), Parus major, Cyanistes cyanus, Periparus ater | 0 | 1 | 1 | 0 | 1 |
| Aves | Corvidae | Corvus frugilegus, Corvus corone, Corvus corax, Corvus monedula (a), Pica pica | 0 | 1 | 1 | 1 | 1 |
| Aves | Garrulus_glandarius | | 0 | 0 | 0 | 1 | 1 |
| Aves | Paridae | Parus major, Cyanistes caeruleus, Periparus ater | 0 | 0 | 0 | 0 | 1 |
| Aves | Regulus_regulus | | 0 | 0 | 0 | 0 | 1 |
| Aves | Sturnus_vulgaris | | 1 | 1 | 1 | 0 | 1 |
| Aves | Sylvia_atricapilla | | 0 | 0 | 0 | 0 | 1 |
| Aves | Bubulcus_ibis | | 0 | 1 | 0 | 0 | 0 |
| Aves | Ardeidae | Ardea alba, Ardea cinerea, Bubulcus ibis | 0 | 0 | 0 | 1 | 1 |
| Aves | Dendrocopos_major | | 0 | 0 | 0 | 0 | 1 |
| Aves | Podiceps_cristatus | | 0 | 0 | 1 | 1 | 1 |
| Aves | Podiceps_nigricollis | | 0 | 1 | 0 | 0 | 0 |
| Aves | Phalacrocorax_carbo_1 | | 1 | 1 | 1 | 1 | 0 |
| Aves | Phalacrocorax_carbo_2 | | 1 | 0 | 1 | 1 | 1 |
| Mammalia | Capreolus_capreolus | | 0 | 0 | 1 | 1 | 1 |
| Mammalia | Vulpes_vulpes | | 0 | 0 | 0 | 1 | 1 |
| Mammalia | Martes_foina | | 0 | 0 | 0 | 0 | 1 |
| Mammalia | Meles_meles | | 0 | 0 | 0 | 0 | 1 |
| Mammalia | Phocidae | Phoca vitulina, Halichoerus grypus | 1 | 1 | 1 | 1 | 0 |
| Mammalia | Delphininae_1 | Tursiops truncatus, Delphinus delphis | 1 | 1 | 1 | 1 | 0 |
| Mammalia | Delphininae_2 | Tursiops truncatus, Delphinus delphis | 1 | 1 | 0 | 0 | 0 |
| Mammalia | Phocoena_phocoena | | 1 | 1 | 0 | 0 | 0 |
| Mammalia | Microtus_sp. | Microtus subterraneus, Microtus agrestis, Microtus arvalis | 0 | 0 | 0 | 0 | 1 |

| | | | | | | | |
|---|---|---|---|---|---|---|---|
| Mammalia | Microtus_arvalis | | 0 | 0 | 0 | 0 | 1 |
| Mammalia | Microtus_subterraneus | | 0 | 0 | 0 | 0 | 1 |
| Mammalia | Myodes_glareolus | | 0 | 0 | 0 | 1 | 1 |
| Mammalia | Ondatra_zibethicus | | 1 | 1 | 1 | 1 | 1 |
| Mammalia | Apodemus_sylvaticus | | 0 | 0 | 1 | 1 | 1 |
| Mammalia | Rattus_norvegicus | | 0 | 1 | 1 | 1 | 1 |
| Mammalia | Myocastor_coypus | | 1 | 1 | 1 | 1 | 1 |
| Mammalia | Sciurus_vulgaris | | 0 | 0 | 0 | 0 | 1 |
| Mammalia | Sorex_sp. | Sorex minutus, Sorex coronatus | 0 | 0 | 0 | 0 | 1 |
| Mammalia | Talpa_europaea | | 0 | 0 | 1 | 0 | 0 |
