## Appendix B for "A Multi-Taxa Approach to Estuarine Biomonitoring: Assessing Vertebrate Biodiversity and Ecological Continuity using Environmental DNA Metabarcoding in the Rance River (Brittany, France)"

***Appendix B*** *"P-values from pairwise permutation tests comparing methods across different biodiversity metrics.*

| Indicator | Station1 | Station2 | p_value |
|---|---|---|---|
| **Taxonomic richness** | ANACONOR | DCE_rance | 1 |
| **Taxonomic richness** | eDNA | DCE_rance | 0.343 |
| **Taxonomic richness** | DCE_rance | ANACONOR | 1 |
| **Taxonomic richness** | eDNA | ANACONOR | 0.678 |
| **Taxonomic richness** | DCE_rance | eDNA | 0.33 |
| **Taxonomic richness** | ANACONOR | eDNA | 0.678 |
| **FRic** | ANACONOR | DCE_rance | 0.321 |
| **FRic** | eDNA | DCE_rance | 1 |
| **FRic** | DCE_rance | ANACONOR | 0.36 |
| **FRic** | eDNA | ANACONOR | 0.656 |
| **FRic** | DCE_rance | eDNA | 1 |
| **FRic** | ANACONOR | eDNA | 0.644 |
| **SES.FRic** | ANACONOR | DCE_rance | 0.313 |
| **SES.FRic** | eDNA | DCE_rance | 1 |

| | | | |
|---|---|---|---|
| **SES.FRic** | DCE_rance | ANACONOR | 0.374 |
| **SES.FRic** | eDNA | ANACONOR | 0.666 |
| **SES.FRic** | DCE_rance | eDNA | 1 |
| **SES.FRic** | ANACONOR | eDNA | 0.636 |
| **PD** | ANACONOR | DCE_rance | 1 |
| **PD** | eDNA | DCE_rance | 0.706 |
| **PD** | DCE_rance | ANACONOR | 1 |
| **PD** | eDNA | ANACONOR | 0.332 |
| **PD** | DCE_rance | eDNA | 0.688 |
| **PD** | ANACONOR | eDNA | 0.333 |
